# Prefrontal metabolite alterations in individuals with posttraumatic stress disorder: a 7T magnetic resonance spectroscopy study

**DOI:** 10.1101/2024.07.16.603137

**Authors:** Meredith A. Reid, Sarah E. Whiteman, Abigail A. Camden, Stephanie M. Jeffirs, Frank W. Weathers

## Abstract

**Background:** Evidence from animal and human studies suggests glutamatergic dysfunction in posttraumatic stress disorder (PTSD). The purpose of this study was to investigate glutamate abnormalities in the dorsolateral prefrontal cortex (DLFPC) of individuals with PTSD using 7T MRS, which has better spectral resolution and signal-to-noise ratio than lower field strengths, thus allowing for better spectral quality and higher sensitivity. We hypothesized that individuals with PTSD would have lower glutamate levels compared to trauma-exposed individuals without PTSD and individuals without trauma exposure. Additionally, we explored potential alterations in other neurometabolites and the relationship between glutamate and psychiatric symptoms.

**Methods:** Individuals with PTSD (n=27), trauma-exposed individuals without PTSD (n=27), and individuals without trauma exposure (n=26) underwent 7T MRS to measure glutamate and other neurometabolites in the left DLPFC. The severities of PTSD, depression, anxiety, and dissociation symptoms were assessed.

**Results:** We found that glutamate was lower in the PTSD and trauma-exposed groups compared to the group without trauma exposure. Furthermore, *N*-acetylaspartate (NAA) was lower and lactate was higher in the PTSD group compared to the group without trauma exposure. Glutamate was negatively correlated with depression symptom severity in the PTSD group. Glutamate was not correlated with PTSD symptom severity.

**Conclusion:** In this first 7T MRS study of PTSD, we observed altered concentrations of glutamate, NAA, and lactate. Our findings provide evidence for multiple possible pathological processes in individuals with PTSD. High-field MRS offers insight into the neurometabolic alterations associated with PTSD and is a powerful tool to probe trauma- and stress-related neurotransmission and metabolism *in vivo*.

## INTRODUCTION

Evidence from animal and human studies points to glutamatergic dysfunction in individuals with posttraumatic stress disorder (PTSD)^1–3^. Fear acquisition, expression, and extinction rely on glutamatergic mechanisms in the amygdala, hippocampus, and prefrontal cortex (PFC)^4, 5^. Studies have shown that stress and corticosterone administration can increase glutamate levels in rats^6–9^, while other studies have demonstrated that the single prolonged stress paradigm^10–12^ and chronic stress^13, 14^ can lower glutamate and glutamine in the PFC of rats. In rats, the single prolonged stress paradigm impairs fear extinction^15^, which is deficient in individuals with PTSD^16^.

Human pharmacological and neuroimaging studies further implicate the glutamate system in the pathophysiology of PTSD. Ketamine, an *N*-methyl-D-aspartate (NMDA) receptor antagonist, reduces PTSD symptom severity^17, 18^. D-cycloserine, a partial NMDA agonist, is somewhat effective in treating PTSD^19–22^. A study using positron emission tomography found higher availability of metabotropic glutamate receptor 5 (mGluR5) in individuals with PTSD, particularly in the orbitofrontal cortex and dorsolateral prefrontal cortex (DLPFC)^23^. Thus, the glutamatergic system is a promising target for drug development^1, 3^.

Magnetic resonance spectroscopy (MRS) is currently the only non-invasive method to measure glutamate *in vivo*. Studies have shown elevated glutamate in the hippocampus^24^ and temporal lobe^25, 26^ of individuals with PTSD, whereas other studies have shown reduced glutamine and Glx (glutamate + glutamine) in the anterior cingulate cortex (ACC)^27, 28^. These discrepant findings could be due to methodological variations, spectral quality differences^29^, or regional heterogeneity. We currently lack PTSD MRS studies of the lateral PFC despite its role in emotion regulation^30–32^ and cognition^33–35^, which are affected in individuals with PTSD^36–39^. Only three PTSD MRS studies examined the DLPFC^40–42^, and only one of those examined Glx, finding no difference between PTSD and control participants^41^. Moreover, most PTSD MRS studies have been conducted at 1.5T or 3T, with none at 7T^43^. MRS at 7T offers better signal-to-noise ratio (SNR) and spectral resolution, which allows for better spectral quality and higher sensitivity to detect more metabolites in a single acquisition^44, 45^. For example, glutamate and glutamine cannot be reliably separated at lower field strengths (≤ 3T) when using conventional MRS sequences because of their overlapping signals, so interpreting the reported differences in Glx (glutamate + glutamine) is difficult.

Similarly, other metabolites, such as gamma-aminobutyric acid (GABA) and glutathione, are challenging to measure with conventional MRS methods at lower field strengths because of their low concentrations and overlap with stronger signals, thus requiring specialized sequences, such as spectral-editing methods, which typically have longer scan times.

In this study, we investigated glutamate alterations in the DLPFC in individuals with PTSD using 7T MRS. Given the prior evidence for reduced Glx in the prefrontal cortex^27, 28^, we hypothesized that individuals with PTSD would have lower glutamate levels compared to trauma-exposed individuals without PTSD and individuals without trauma exposure. Additionally, we explored the relationship between glutamate and psychiatric symptoms as well as potential alterations in other neurometabolites.

## METHODS

### Participants

The Auburn University Institutional Review Board approved this study. All participants provided written informed consent. The study sample comprised 80 people, ages 19 to 55, recruited from the local community (n=65) and from undergraduate students enrolled in psychology courses at Auburn University (n=15; Supplementary Methods).

We used the Life Events Checklist for *DSM-5* extended version to determine *DSM-5* Criterion A status based on participants’ self-identified worst event (i.e., index event) and the PTSD Checklist for *DSM-5* (PCL-5) to measure PTSD symptoms^46^ (Supplementary Methods). Participants also completed the Beck Depression Inventory (BDI-II), Beck Anxiety Inventory (BAI), and Multiscale Dissociation Inventory (MDI).

Psychiatric comorbidities were obtained from the Mini International Neuropsychiatric Interview for *DSM-5* (version 7.0.0; n=16) or from participants’ self-reported psychiatric history (n=64). Information about substance use was obtained from n=64 participants using the *DSM-5* Self-Rated Level 1 Cross-Cutting Symptom Measure^47^ (Supplementary Methods).

Participants were included in the PTSD group if they met all the following criteria: (1) their self-identified worst event (i.e., index event) was a *DSM-5* Criterion A event (Supplementary Methods); (2) they met *DSM-5* diagnostic criteria using PCL-5 scoring rules (Supplementary Methods); and (3) they had a PCL-5 total score of at least 30 to increase confidence in the provisional PTSD diagnosis. PCL-5 total scores of 31-33 and above have been shown to indicate probable PTSD, although studies have used cutoff scores ranging from 23 to 49, and a universal cutoff score has not been established^48^. Participants were included in the trauma-exposed (TE) group if they met all the following criteria: (1) their index event was a Criterion A event (Supplementary Methods); (2) they either did not meet full diagnostic criteria for PTSD based on the PCL-5 or had a PCL-5 total score of less than 30 (Supplementary Methods); and (3) their BDI and BAI scores were moderate or below (BDI≤28 and BAI≤25). Participants were included in the no trauma exposure (NT) group if they met all the following criteria: (1) their index event was not a Criterion A event (Supplementary Methods); (2) they did not endorse directly experiencing any other events (Supplementary Methods); and (3) their BDI and BAI scores were moderate or below (BDI≤28 and BAI≤25). Exclusion criteria for the study were neurological disorders, history of concussion with loss of consciousness, alcohol use disorder, substance use disorder, psychosis and schizophrenia spectrum disorders, bipolar and related disorders, obsessive-compulsive disorder, personality disorders, and autism spectrum disorder.

### MR acquisition and analysis

Data were acquired on a Siemens 7T MAGNETOM scanner (Supplementary Methods; Table S1). Three-dimensional structural images were acquired for anatomical reference and MRS voxel segmentation (Supplementary Methods). The DLPFC voxel (25×25×25 mm^3^) was oriented parallel to the cortical surface when viewed in the sagittal and axial planes^49^ (Figure 1A). A large voxel was chosen because these data were acquired as part of a study that included functional MRS for which a larger voxel is advantageous for SNR when measuring dynamic changes in metabolites during shorter acquisitions. Participants were instructed to fixate on a crosshair during the MRS scan. Spectra were acquired using an ultra-short TE STEAM sequence^49–52^ (Supplementary Methods; Table S1).

**Figure 1.**
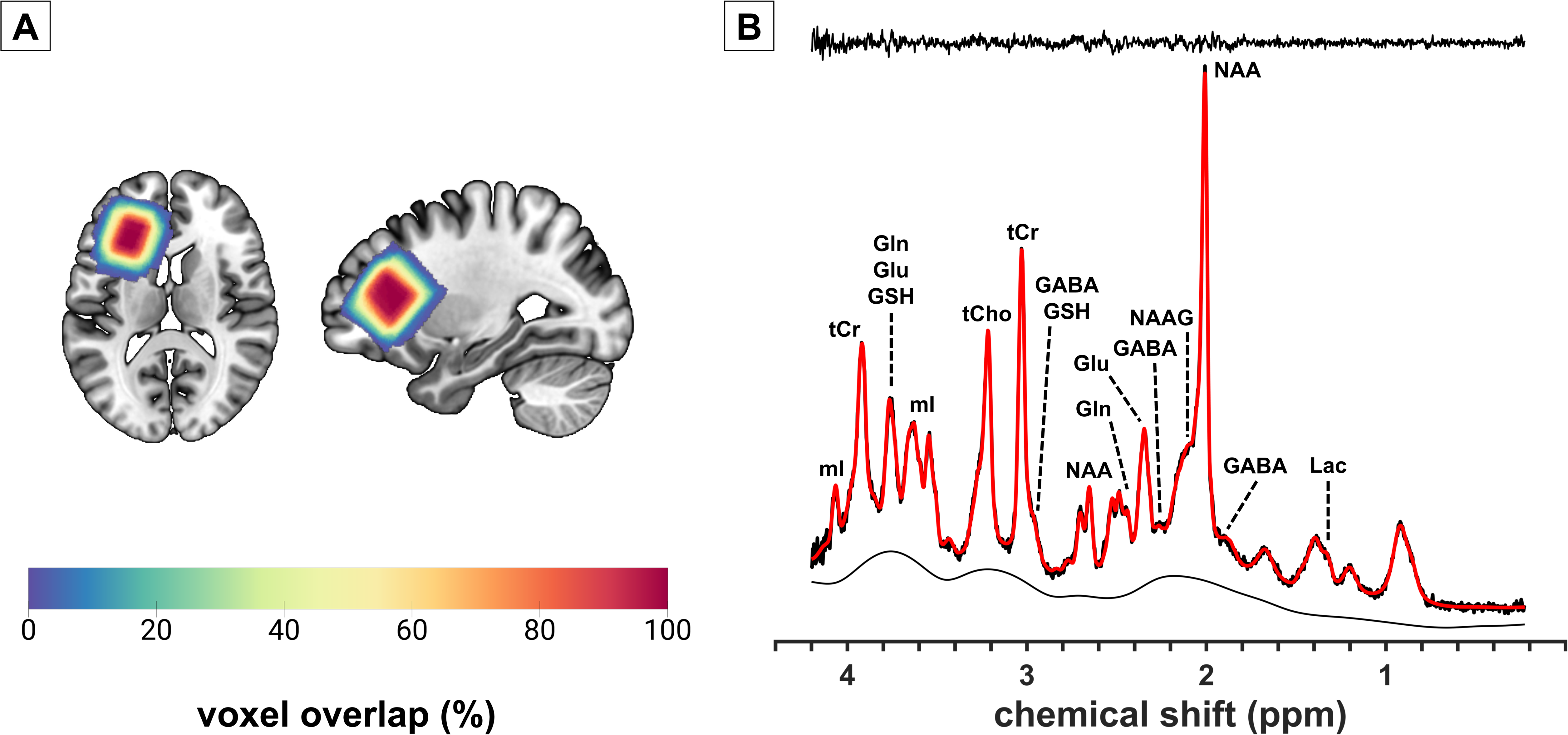
(A) MRS voxel position in the left dorsolateral prefrontal cortex. The color scale represents the overlap in voxel position across participants. (B) Representative spectrum from the dorsolateral prefrontal cortex. Red is the LCModel fit overlaid on the acquired spectrum. The residual signal (the difference between the spectrum and fit) is shown above the spectrum, and the background signal is shown below the spectrum. Group-averaged spectra are shown in Figure S1. Abbreviations: GABA, gamma-aminobutyric acid; Gln, glutamine; Glu, glutamate; GSH, glutathione; Lac, lactate; mI, myo-inositol; NAA, *N*-acetylaspartate; NAAG, *N*-acetylaspartylglutamate; ppm, parts per million; tCho, total choline (glycerophosphocholine + phosphocholine); tCr, total creatine (creatine + phosphocreatine); tNAA, total NAA (NAA + NAAG).

Spectra were processed with Osprey^53^ (version 2.4.0) and LCModel^54^ (version 6.3-1N implemented in Osprey; Supplementary Methods; Table S1). Osprey segmented the tissue within the MRS voxel into gray matter, white matter, and cerebrospinal fluid using SPM12. Osprey calculated the tissue- and relaxation-corrected molal concentration estimates (mmol/kg of tissue water) for each metabolite^53, 55^. Osprey also calculated the SNR of creatine, the linewidth of creatine, and the linewidth of the unsuppressed water signal.

Spectra were visually inspected for quality and fit. Exclusion criteria were poor fit, poor water suppression, water linewidth greater than 30 Hz, or creatine linewidth greater than 30 Hz (Table S2). Individual metabolite concentrations were excluded from statistical analysis if the Cramer-Rao lower bounds (CRLB) were greater than 20%, except for the smaller signals of lactate and *N*-acetylaspartylglutamate (NAAG) for which the threshold was 30% (Table S2).

### Statistical analysis

Statistical analyses were conducted in R/RStudio (Supplementary Methods). Statistical significance was set at two-tailed p<0.05. ANOVA was used for group comparisons of age, BDI total score, BAI total score, MDI total score, CRLB, SNR, linewidth, and tissue fractions. Post hoc pairwise t-tests with Tukey-adjusted p-values were conducted if the omnibus test was significant. PCL-5 total score and years since trauma exposure were compared between the PTSD and TE groups using t-tests. Chi-square tests were used to compare sex, trauma in childhood/adolescence or adulthood, and the presence of comorbid psychiatric disorders between groups. Fisher’s exact test was used to compare race and ethnicity, index trauma category (natural disaster, assault, etc.), category of index trauma exposure (direct, witnessed, learned about, or job), and psychotropic medication status (currently medicated or unmedicated). Post hoc pairwise comparisons with Bonferroni-adjusted p-values were conducted if the omnibus test was significant.

For statistical analysis of metabolite concentrations, the median absolute deviation was used to identify outliers. Individual metabolite concentrations were excluded if they differed from the median value by more than three times the median absolute deviation^56^ (Table S2). Metabolites were compared between groups using ANCOVA controlling for age and sex^57^. Statistical significance was set at Bonferroni-corrected p<0.0036 (0.05/14 metabolites). Post hoc pairwise t-tests with Tukey-adjusted p-values were conducted if the omnibus test was significant. Pearson correlation coefficients were used to explore the association between glutamate and scores on the PCL-5, BDI, BAI, and MDI. Statistical significance was set at Bonferroni-corrected p<0.0125 (0.05/4 symptom scales). Correlations were compared between groups using the Fisher r-to-z transform.

In secondary analyses, the group comparisons were repeated after performing alpha correction^58^ for glutamate, glutamine, Glx, and GABA and after including the gray matter fraction (gray matter / (gray matter + white matter)) as a covariate for all metabolite group comparisons. Alpha correction accounts for concentration differences in gray matter and white matter and normalizes the concentrations to a standard voxel composition based on the group-averaged composition^58–60^. Osprey implements alpha correction for glutamate, glutamine, Glx, and GABA with an assumed alpha of 0.5, which represents a 2:1 concentration ratio between gray matter and white matter. Alpha correction is preferred over the tissue covariate approach because the latter reduces statistical power^59^ and complicates the interpretation because the tissue fraction is related to both the dependent (metabolite) and independent (group, age) variables^61^.

## RESULTS

### Demographics and clinical characteristics

The sample included 27 PTSD, 27 TE, and 26 NT participants (Table 1; Table S3). Age, sex, race, and ethnicity were not significantly different between the groups. PCL-5 was significantly higher in the PTSD group compared to the TE group (p<0.001). BDI was significantly higher in the PTSD group compared to the TE group (p_Tukey_<0.001) and the NT group (p_Tukey_<0.001). BAI was significantly higher in the PTSD group compared to the TE group (p_Tukey_<0.001) and the NT group (p_Tukey_<0.001) and significantly higher in the TE group compared to the NT group (p_Tukey_=0.02). MDI was significantly higher in the PTSD group compared to the TE group (p_Tukey_<0.001) and the NT group (p_Tukey_<0.001). Index trauma category, category of trauma exposure, and the time since trauma exposure were not significantly different between the PTSD and TE groups (Table 1). The number of participants whose index event occurred during childhood or adolescence compared to adulthood was not significantly different between the PTSD and TE groups. The PTSD group had significantly more participants with psychiatric comorbidities than the NT group (p_Bonferroni_<0.001). The PTSD group had significantly more participants taking psychotropic medications than the NT group (pBonferroni=0.04).

**Table 1.** Demographics and clinical measures.

|  | **PTSD** | **TE** | **NT** | **Statistic** | **p-value** |
| --- | --- | --- | --- | --- | --- |
| Sample size | 27 | 27 | 26 |  |  |
| Age, Mean (SD) | 27.85 (8.32) | 31.56 (9.20) | 29.08 (10.97) | $F_{2,77} = 1.06$ | 0.35 |
| Sex, n | | | | $\chi^2_2 = 4.01$ | 0.14 |
| Female | 21 | 14 | 16 |  |  |
| Male | 6 | 13 | 10 |  |  |
| Race, n |  |  |  | Fisher's exact | 0.55 |
| Asian | 1 | 1 | 4 |  |  |
| Multiracial | 2 | 2 | 1 |  |  |
| White | 24 | 24 | 21 |  |  |
| Ethnicity, n |  |  |  | Fisher's exact | 0.87 |
| Hispanic or Latino | 2 | 3 | 1 |  |  |
| Non-Hispanic or Non-Latino | 25 | 24 | 25 |  |  |
| BDI, Mean (SD) | 26.37 (10.16)<sup>a,b</sup> | 7.96 (6.14) | 4.81 (6.00) | $F_{2,77} = 61.24$ | < 0.001 |
| BAI, Mean (SD) | 25.41 (9.94)<sup>a,b</sup> | 8.15 (6.83)<sup>c</sup> | 2.65 (3.26) | $F_{2,77} = 71.58$ | < 0.001 |
| MDI, Mean (SD) | 72.93 (17.47)<sup>a,b</sup> | 43.00 (13.13) | 36.92 (7.88) | $F_{2,77} = 54.74$ | < 0.001 |
| PCL-5, Mean (SD) | 50.93 (11.59) | 11.52 (9.63) | --- | $t_{52} = 13.59$ | < 0.001 |
| Index trauma, n |  |  |  | Fisher's exact | 0.41 |
| Natural disaster | 0 | 1 | --- |  |  |
| Fire or explosion | 1 | 1 | --- |  |  |
| Transportation accident | 3 | 3 | --- |  |  |
| Serious accident | 0 | 4 | --- |  |  |
| Physical assault | 5 | 2 | --- |  |  |
| Assault with a weapon | 4 | 3 | --- |  |  |
| Sexual assault | 6 | 2 | --- |  |  |
| Other unwanted or uncomfortable sexual experience | 1 | 1 | --- |  |  |
| Combat or exposure to a warzone | 1 | 3 | --- |  |  |
| Life-threatening illness or injury | 2 | 1 | --- |  |  |
| Sudden violent death | 4 | 6 | --- |  |  |
| Exposure, n |  |  |  | Fisher's exact | 0.56 |
| Direct | 19 | 16 | --- |  |  |
| Witnessed | 3 | 3 | --- |  |  |
| Learned about | 3 | 7 | --- |  |  |
| Job | 2 | 1 | --- |  |  |
| Years since trauma exposure, Mean (SD)<sup>d</sup> | 6.68 (5.94) | 10.33 (10.85) | --- | $t_{51} = 1.52$ | 0.13 |
| Trauma in childhood/adolescence or adulthood, n | | | | $\chi^2_1 = 0.67$ | 0.41 |
| Childhood or adolescence | 16 | 12 | --- |  |  |
| Adulthood | 11 | 15 | --- |  |  |
| Comorbid disorders, yes/no<sup>e</sup> | 21/6<sup>b</sup> | 14/13 | 6/20 | $\chi^2_2 = 15.87$ | < 0.001 |
| Anxiety | 19 | 6 | 0 |  |  |
| ADHD | 10 | 3 | 0 |  |  |
| Depression/MDD | 16 | 5 | 6 |  |  |
| Eating | 5 | 1 | 0 |  |  |
| Panic | 5 | 1 | 0 |  |  |
| Phobia | 1 | 1 | 0 |  |  |
| PMDD | 0 | 1 | 0 |  |  |
| Sleep | 5 | 2 | 0 |  |  |
| Somatoform | 1 | 0 | 0 |  |  |
| Psychotropic medications, yes/no<sup>f</sup> | 11/16<sup>b</sup> | 5/22 | 2/24 | Fisher's exact | 0.02 |
| Alprazolam | 1 | 0 | 0 |  |  |
| Amphetamine | 2 | 0 | 0 |  |  |
| Bupropion | 2 | 1 | 0 |  |  |
| Buspirone | 3 | 1 | 0 |  |  |
| Duloxetine | 1 | 0 | 0 |  |  |
| Escitalopram | 1 | 0 | 0 |  |  |
| Fluoxetine | 0 | 1 | 0 |  |  |
| Lamotrigine | 2 | 0 | 0 |  |  |
| Lisdexamfetamine | 1 | 2 | 0 |  |  |
| Methylphenidate | 1 | 0 | 0 |  |  |
| Paroxetine | 0 | 0 | 1 |  |  |
| Quetiapine | 1 | 0 | 0 |  |  |
| Sertraline | 2 | 0 | 0 |  |  |
| Trazodone | 2 | 0 | 0 |  |  |
| Venlafaxine | 1 | 0 | 1 |  |  |

Abbreviations: ADHD, attention-deficit/hyperactivity disorder; BAI, Beck Anxiety Inventory; BDI, Beck Depression Inventory; F, female; M, male; MDD, major depressive disorder; MDI, Multiscale Dissociation Inventory; NT, no trauma exposure; PCL-5, PTSD Checklist for *DSM-5*; PMDD, premenstrual dysphoric disorder; PTSD, posttraumatic stress disorder; TE, trauma-exposed without PTSD
<sup>a</sup>Significant difference between PTSD and TE (see Table S3).
<sup>b</sup>Significant difference between PTSD and NT (see Table S3).
<sup>c</sup>Significant difference between TE and NT (see Table S3).
<sup>d</sup>Information was not available for 1 TE. The participant indicated that the index event occurred in adulthood but did not provide sufficient details to determine a more precise time since trauma exposure.
<sup>e</sup>Some participants had more than one comorbid disorder.
<sup>f</sup>Some participants were taking more than one psychotropic medication.

### Glutamate and other metabolites

Three spectra were excluded from the analysis due to poor water suppression (1 PTSD) and Cr linewidth greater than 30 Hz (2 NT; Table S2). Voxel overlap across participants and a representative spectrum are shown in Figure 1. Group-averaged spectra are shown in Figure S1. Metabolite concentrations, spectral quality metrics, and tissue fractions are provided in Table 2. SNR and linewidths did not significantly differ between the groups. Gray and white matter tissue fractions were significantly different between the groups (Table 2; Table S4). Gray matter was lower in the TE group compared to the NT group (p_Tukey_=0.049), and white matter was higher in the PTSD group (p_Tukey_=0.049) and the TE group (p_Tukey_=0.04) compared to the NT group.

**Table 2.**
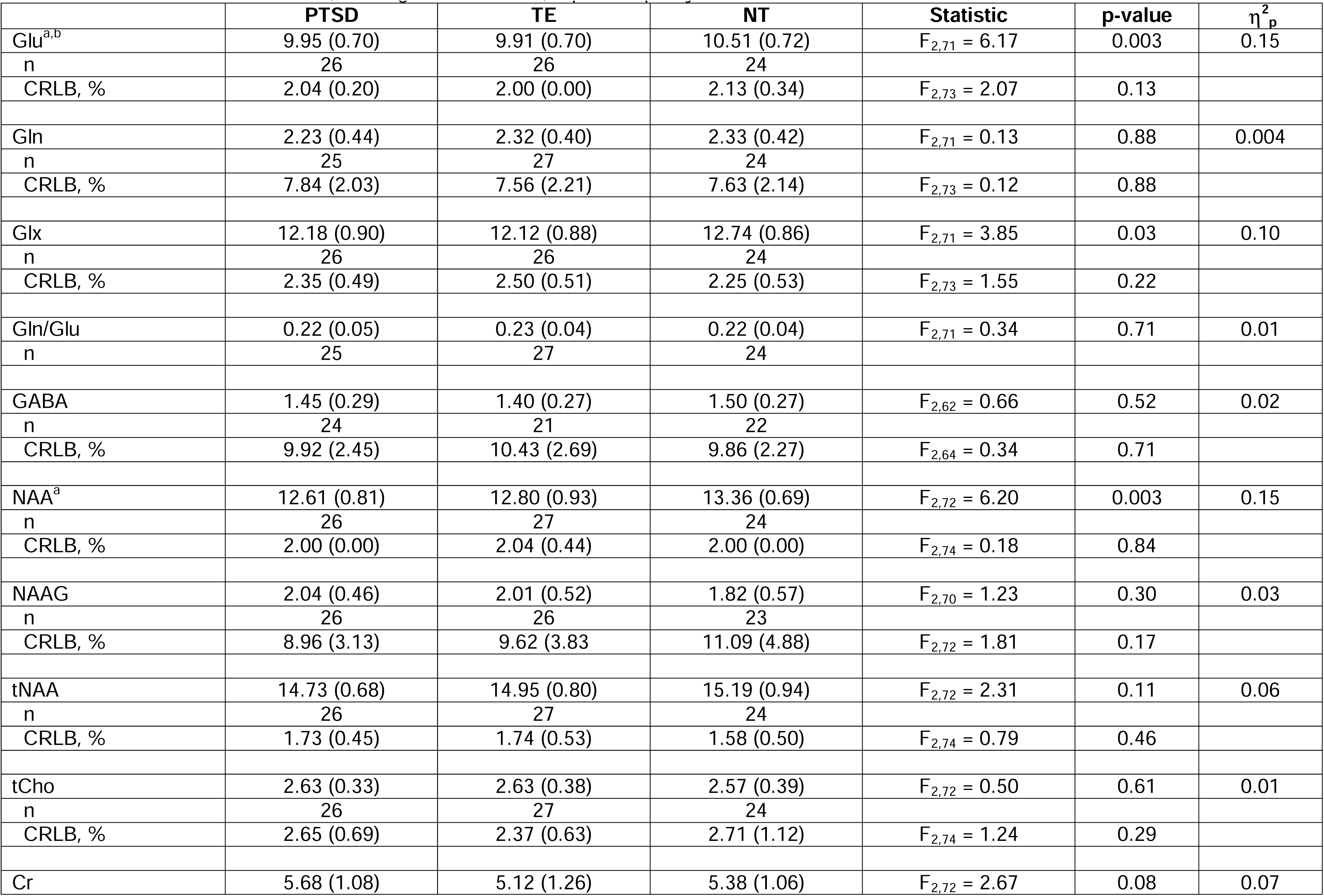
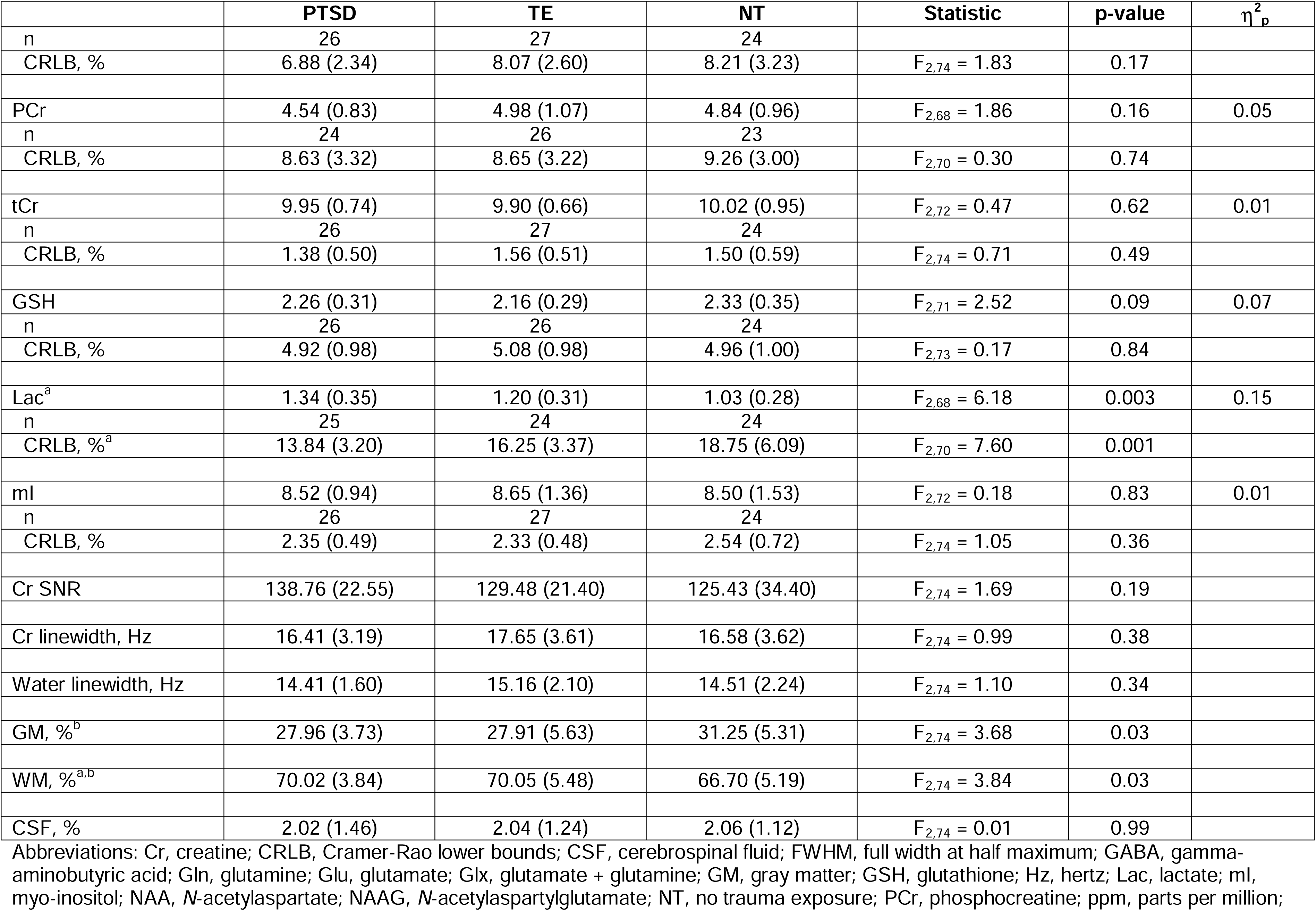
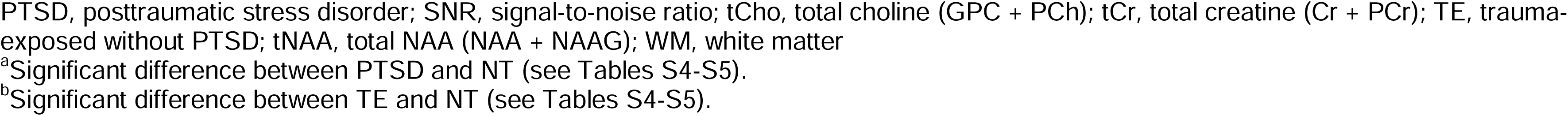
Metabolite concentrations (mmol/kg of tissue water), spectral quality, and tissue fractions.

|  | **PTSD** | **TE** | **NT** | **Statistic** | **p-value** | **$\eta^2_p$** |
| --- | --- | --- | --- | --- | --- | --- |
| Glu<sup>a,b</sup> | 9.95 (0.70) | 9.91 (0.70) | 10.51 (0.72) | $F_{2,71} = 6.17$ | 0.003 | 0.15 |
| n | 26 | 26 | 24 |  |  |  |
| CRLB, % | 2.04 (0.20) | 2.00 (0.00) | 2.13 (0.34) | $F_{2,73} = 2.07$ | 0.13 | |
| Gln | 2.23 (0.44) | 2.32 (0.40) | 2.33 (0.42) | $F_{2,71} = 0.13$ | 0.88 | 0.004 |
| n | 25 | 27 | 24 |  |  |  |
| CRLB, % | 7.84 (2.03) | 7.56 (2.21) | 7.63 (2.14) | $F_{2,73} = 0.12$ | 0.88 | |
| Glx | 12.18 (0.90) | 12.12 (0.88) | 12.74 (0.86) | $F_{2,71} = 3.85$ | 0.03 | 0.10 |
| n | 26 | 26 | 24 |  |  |  |
| CRLB, % | 2.35 (0.49) | 2.50 (0.51) | 2.25 (0.53) | $F_{2,73} = 1.55$ | 0.22 | |
| Gln/Glu | 0.22 (0.05) | 0.23 (0.04) | 0.22 (0.04) | $F_{2,71} = 0.34$ | 0.71 | 0.01 |
| n | 25 | 27 | 24 |  |  |  |
| GABA | 1.45 (0.29) | 1.40 (0.27) | 1.50 (0.27) | $F_{2,62} = 0.66$ | 0.52 | 0.02 |
| n | 24 | 21 | 22 |  |  |  |
| CRLB, % | 9.92 (2.45) | 10.43 (2.69) | 9.86 (2.27) | $F_{2,64} = 0.34$ | 0.71 | |
| NAA<sup>a</sup> | 12.61 (0.81) | 12.80 (0.93) | 13.36 (0.69) | $F_{2,72} = 6.20$ | 0.003 | 0.15 |
| n | 26 | 27 | 24 |  |  |  |
| CRLB, % | 2.00 (0.00) | 2.04 (0.44) | 2.00 (0.00) | $F_{2,74} = 0.18$ | 0.84 | |
| NAAG | 2.04 (0.46) | 2.01 (0.52) | 1.82 (0.57) | $F_{2,70} = 1.23$ | 0.30 | 0.03 |
| n | 26 | 26 | 23 |  |  |  |
| CRLB, % | 8.96 (3.13) | 9.62 (3.83) | 11.09 (4.88) | $F_{2,72} = 1.81$ | 0.17 | |
| tNAA | 14.73 (0.68) | 14.95 (0.80) | 15.19 (0.94) | $F_{2,72} = 2.31$ | 0.11 | 0.06 |
| n | 26 | 27 | 24 |  |  |  |
| CRLB, % | 1.73 (0.45) | 1.74 (0.53) | 1.58 (0.50) | $F_{2,74} = 0.79$ | 0.46 | |
| tCho | 2.63 (0.33) | 2.63 (0.38) | 2.57 (0.39) | $F_{2,72} = 0.50$ | 0.61 | 0.01 |
| n | 26 | 27 | 24 |  |  |  |
| CRLB, % | 2.65 (0.69) | 2.37 (0.63) | 2.71 (1.12) | $F_{2,74} = 1.24$ | 0.29 | |
| Cr | 5.68 (1.08) | 5.12 (1.26) | 5.38 (1.06) | $F_{2,72} = 2.67$ | 0.08 | 0.07 |

| | **PTSD** | **TE** | **NT** | **Statistic** | **p-value** | $\eta^2_p$ |
| --- | --- | --- | --- | --- | --- | --- |
| n | 26 | 27 | 24 |  |  |  |
| CRLB, % | 6.88 (2.34) | 8.07 (2.60) | 8.21 (3.23) | $F_{2,74} = 1.83$ | 0.17 | |
| PCr | 4.54 (0.83) | 4.98 (1.07) | 4.84 (0.96) | $F_{2,68} = 1.86$ | 0.16 | 0.05 |
| n | 24 | 26 | 23 |  |  |  |
| CRLB, % | 8.63 (3.32) | 8.65 (3.22) | 9.26 (3.00) | $F_{2,70} = 0.30$ | 0.74 | |
| tCr | 9.95 (0.74) | 9.90 (0.66) | 10.02 (0.95) | $F_{2,72} = 0.47$ | 0.62 | 0.01 |
| n | 26 | 27 | 24 |  |  |  |
| CRLB, % | 1.38 (0.50) | 1.56 (0.51) | 1.50 (0.59) | $F_{2,74} = 0.71$ | 0.49 | |
| GSH | 2.26 (0.31) | 2.16 (0.29) | 2.33 (0.35) | $F_{2,71} = 2.52$ | 0.09 | 0.07 |
| n | 26 | 26 | 24 |  |  |  |
| CRLB, % | 4.92 (0.98) | 5.08 (0.98) | 4.96 (1.00) | $F_{2,73} = 0.17$ | 0.84 | |
| Lac<sup>a</sup> | 1.34 (0.35) | 1.20 (0.31) | 1.03 (0.28) | $F_{2,68} = 6.18$ | 0.003 | 0.15 |
| n | 25 | 24 | 24 |  |  |  |
| CRLB, %<sup>a</sup> | 13.84 (3.20) | 16.25 (3.37) | 18.75 (6.09) | $F_{2,70} = 7.60$ | 0.001 | |
| ml | 8.52 (0.94) | 8.65 (1.36) | 8.50 (1.53) | $F_{2,72} = 0.18$ | 0.83 | 0.01 |
| n | 26 | 27 | 24 |  |  |  |
| CRLB, % | 2.35 (0.49) | 2.33 (0.48) | 2.54 (0.72) | $F_{2,74} = 1.05$ | 0.36 | |
| Cr SNR | 138.76 (22.55) | 129.48 (21.40) | 125.43 (34.40) | $F_{2,74} = 1.69$ | 0.19 | |
| Cr linewidth, Hz | 16.41 (3.19) | 17.65 (3.61) | 16.58 (3.62) | $F_{2,74} = 0.99$ | 0.38 | |
| Water linewidth, Hz | 14.41 (1.60) | 15.16 (2.10) | 14.51 (2.24) | $F_{2,74} = 1.10$ | 0.34 | |
| GM, %<sup>b</sup> | 27.96 (3.73) | 27.91 (5.63) | 31.25 (5.31) | $F_{2,74} = 3.68$ | 0.03 | |
| WM, %<sup>a,b</sup> | 70.02 (3.84) | 70.05 (5.48) | 66.70 (5.19) | $F_{2,74} = 3.84$ | 0.03 | |
| CSF, % | 2.02 (1.46) | 2.04 (1.24) | 2.06 (1.12) | $F_{2,74} = 0.01$ | 0.99 | |

Abbreviations: Cr, creatine; CRLB, Cramer-Rao lower bounds; CSF, cerebrospinal fluid; FWHM, full width at half maximum; GABA, gamma-aminobutyric acid; Gln, glutamine; Glu, glutamate; Glx, glutamate + glutamine; GM, gray matter; GSH, glutathione; Hz, hertz; Lac, lactate; ml, myo-inositol; NAA, *N*-acetylaspartate; NAAG, *N*-acetylaspartylglutamate; NT, no trauma exposure; PCr, phosphocreatine; ppm, parts per million;
PTSD, posttraumatic stress disorder; SNR, signal-to-noise ratio; tCho, total choline (GPC + PCh); tCr, total creatine (Cr + PCr); TE, trauma-exposed without PTSD; tNAA, total NAA (NAA + NAAG); WM, white matter
<sup>a</sup>Significant difference between PTSD and NT (see Tables S4-S5).
<sup>b</sup>Significant difference between TE and NT (see Tables S4-S5).

There was a significant group difference in glutamate after controlling for age and sex (Figure 2A; Table 2; Table S5). Glutamate was significantly lower in the PTSD group compared to the NT group (p_Tukey_=0.005, Cohen’s d=0.92) and significantly lower in the TE group compared to the NT group (p_Tukey_=0.02, Cohen’s d=0.80). There was a significant group difference in *N*-acetylaspartate (NAA) after controlling for age and sex (Figure 2B; Table 2; Table S5). NAA was significantly lower in the PTSD group compared to the NT group (p_Tukey_=0.002, Cohen’s d=0.99). There was a significant group difference in lactate after controlling for age and sex (Figure 2C; Table 2; Table S5). Lactate was significantly higher in the PTSD group compared to the NT group (p_Tukey_=0.002, Cohen’s d=1.01). There were non-significant trends for lower Glx and glutathione in the PTSD and TE groups compared to the NT group, and no significant group differences for the other metabolites (Table 2; Figure S2). In a secondary analysis, the group difference in the glutamine/glutamate ratio was not significant (Table 2; Figure S2).

**Figure 2.**
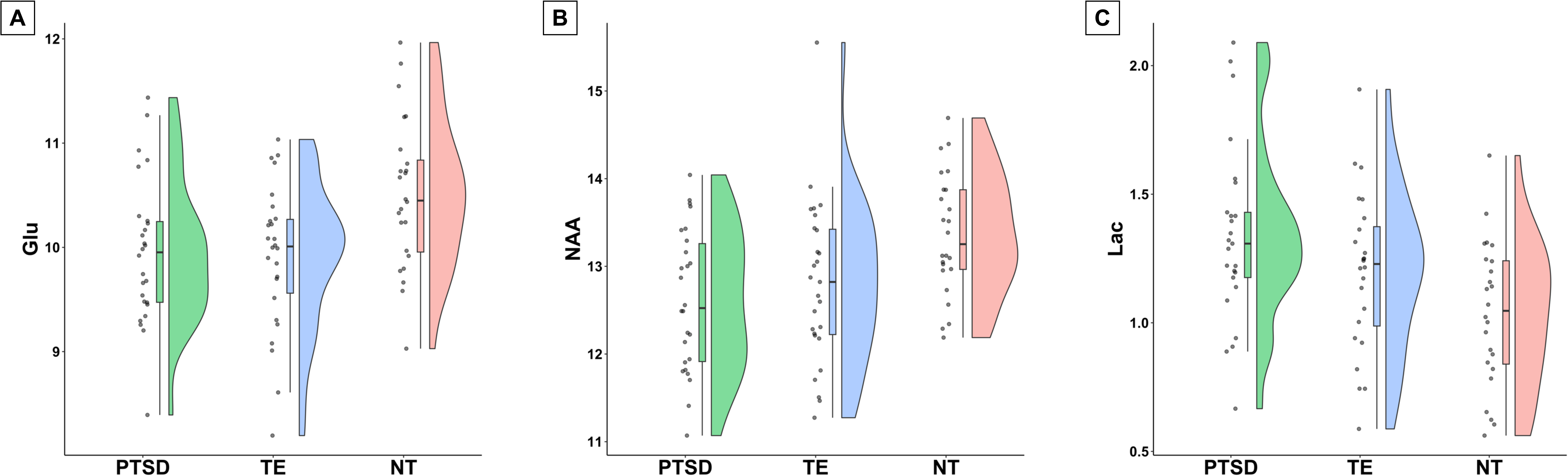
Concentrations (mmol/kg of tissue water) of (A) glutamate, (B) *N*- acetylaspartate, and (C) lactate. Abbreviations: Glu, glutamate; Lac, lactate; NAA, *N*- acetylaspartate; NT, no trauma exposure; PTSD, posttraumatic stress disorder; TE, trauma-exposed without PTSD.

Following alpha correction, the group difference remained significant for glutamate (F_2,71_=6.19, p=0.003, η^2^ =0.15; Table S6). When including the tissue fraction as a covariate, the group differences in glutamate (F_2,70_=2.76, p=0.07, η^2^ =0.07; Table S7), NAA (F_2,71_=2.98, p=0.06, η^2^ =0.08; Table S7), and lactate (F_2,67_=4.25, p=0.02, η^2^ =0.11; Table S7) were not significant at the Bonferroni-corrected p<0.0036.

In an exploratory analysis, we compared glutamate, NAA, and lactate between adulthood trauma and childhood/adolescence trauma in the combined sample of PTSD and TE participants. The difference was not statistically significant for glutamate, NAA, and lactate after covarying for age and sex (Table S8; Figure S3). We also assessed the robustness of our results after accounting for potential confounding factors, such as medication status and substance use, and the results were similar (Supplementary Results; Tables S9-S14). Group comparisons without the removal of metabolite outliers (see Methods and Table S2) are provided in Table S15 and Figure S4.

### Glutamate correlation with symptoms

Glutamate was negatively correlated with BDI scores in the combined sample (r_74_= -0.23, p=0.046), and this association was driven by the PTSD group (PTSD: r_24_= - 0.47, p=0.01; TE: r_24_=0.17, p=0.42; NT: r_22_=0.19, p=0.36; PTSD vs. TE: z=2.31, p=0.02; PTSD vs. NT: z=2.33, p=0.02; TE vs. NT: z=0.07, p=0.95; Figure 3A). There was a non-significant negative trend between glutamate and MDI scores in the PTSD group (PTSD: r_24_= -0.36, p=0.07; TE: r_24_=0.03, p=0.87; PTSD+TE: r_50_= -0.11, p=0.44; PTSD vs. TE: z=1.39, p=0.17; Figure 3B). Glutamate was not significantly correlated with PCL- 5 scores (PTSD: r_24_= -0.10, p=0.63; TE: r_24_=0.09, p=0.65; PTSD+TE: r_50_=0.02, p=0.89; PTSD vs. TE: z=0.66, p=0.51; Figure S5) or BAI scores (combined sample: r_74_= -0.15, p=0.21; PTSD: r_24_=0.06, p=0.78; TE: r_24_=0.07, p=0.74; NT: r_22_=0.16, p=0.47; PTSD vs. TE: z=0.04, p=0.97; PTSD vs. NT: z=0.34, p=0.73; TE vs. NT: z=0.31, p=0.76; Figure S6).

**Figure 3.**
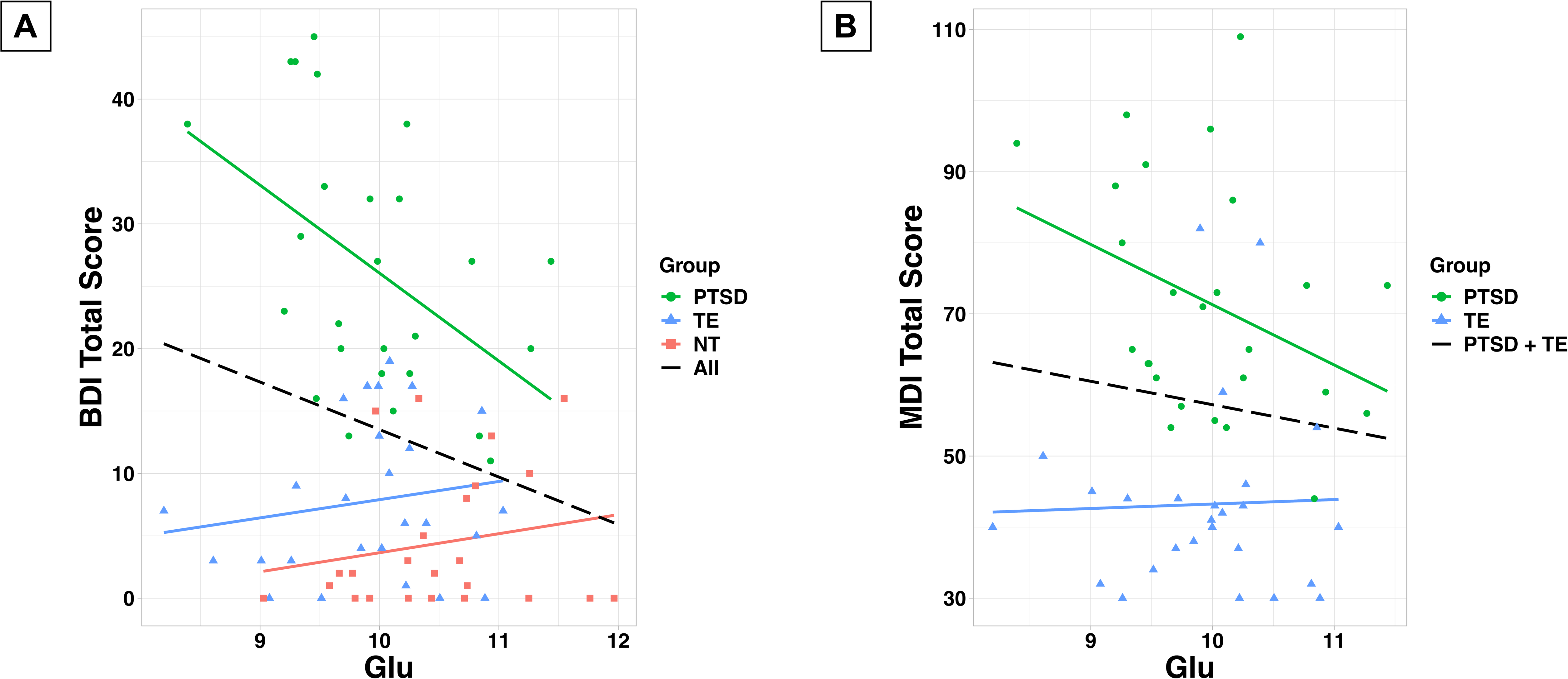
(A) Correlation between glutamate concentration (mmol/kg of tissue water) and Beck Depression Inventory (BDI) total score. (B) Correlation between glutamate concentration (mmol/kg of tissue water) and Multiscale Dissociation Inventory (MDI) total score.

## DISCUSSION

We investigated glutamate and other neurometabolites in the DLPFC of individuals with PTSD using 7T proton MRS. We observed that individuals with PTSD had lower glutamate, lower NAA, and elevated lactate compared to individuals without trauma exposure. Glutamate was also reduced in trauma-exposed individuals without PTSD compared to those without trauma exposure. Glutamate was negatively correlated with depression symptom severity in the PTSD group.

We observed lower glutamate in both the PTSD and trauma-exposed groups compared to the group without trauma exposure. This finding is similar to studies that found lower Glx in the dorsal ACC and medial PFC among trauma survivors, including individuals with PTSD, panic disorder, major depressive disorder, and generalized anxiety disorder^62, 63^. Other studies specifically focused on individuals with PTSD have reported lower Glx^27^ and glutamine^28^ in the ACC, elevated glutamate in the hippocampus^24^ and temporal lobe^25, 26^, and no differences in glutamate, glutamine, or Glx in the DLPFC^41^ and posterior cingulate gyrus^64^.

In contrast to other studies^24–26, 65^, we found no association between glutamate and PTSD symptom severity. However, we found that glutamate was negatively correlated with depression symptom severity in the PTSD group, suggesting lower glutamate may be associated more with depressive symptomatology than with PTSD diagnosis. To our knowledge, no other PTSD MRS studies have reported an association between glutamate and depression symptom severity^43^. Notably, although both the PTSD and trauma-exposed groups had lower glutamate compared to the group without trauma exposure, we observed the association between glutamate and depression symptom severity only in the PTSD group. While our sample included participants with both single-event and repeated trauma exposure, most participants, especially those in the PTSD group, experienced repeated trauma exposure. Specifically, 21/27 of the participants in the PTSD group and 11/27 of the participants in the trauma-exposed group reported experiencing their index events more than once. This repeated exposure may be indicative of higher levels of trauma exposure and chronic stress. Given the relationship between chronic stress, depression, and altered glutamate^66^, it is not surprising that glutamate was correlated with depression severity in the PTSD group. The range of depression severity was smaller in the trauma-exposed group, which may explain the lack of correlation in this group, so future studies should examine a wider range of depression severities in both trauma-exposed and non-trauma-exposed individuals. We also observed a trend for lower glutamate associated with greater dissociation symptoms in the PTSD group. In a recent study, lower GABA in the medial PFC was associated with higher trait dissociation in trauma-exposed individuals with and without PTSD^67^. Our findings collectively underscore the need for more studies of the relationship between PFC neurobiology and the symptomatology of depression and dissociation.

Our findings and previous studies suggest that glutamatergic dysfunction in individuals with PTSD could be due to disrupted glucocorticoid signaling^1^, astrocyte dysfunction^1, 68, 69^, or increased inflammation^70^. These factors may impair glutamatergic transmission and metabolism in the PFC^1^. Studies have shown that acute stress and corticosterone administration increased extracellular glutamate in rats^1, 6–8^, whereas chronic stress lowered glutamate^13, 14^. The single prolonged stress paradigm also resulted in lower glutamate and glutamine in the rat PFC measured seven days after removing the stressor^10–12^, demonstrating that acute stress can have persistent effects on glutamate signaling^69^. Stress negatively impacts astrocytes^1, 68^, which are crucial for glutamate clearance and metabolism. Chronic unpredictable stress was associated with reduced glutamate-glutamine cycling in the rat PFC^71^. Although we found no differences in the glutamine-glutamate ratio, a recent study using ^13^C-MRS to examine PFC glutamate neurotransmission *in vivo*^72^ found that individuals with PTSD had lower energy-per-cycle, which is the ratio of the rate of neuronal oxidative energy production (V_TCA_) to the rate of glutamate-glutamine cycling (V_Cycle_)^72^. Astrocyte dysfunction and disrupted uptake of glutamate could increase extracellular glutamate, leading to excitotoxicity^1, 3^ and subsequently inducing morphological changes. In rodents, acute and chronic stress^73–76^ and corticosterone administration^77, 78^ have been shown to cause structural abnormalities in the PFC and hippocampus, including reduced dendritic branching and loss of spines^79^. Chronic stress and decreased spine density were correlated with deficits in working memory and other executive functions^79, 80^, which are present in individuals with PTSD^36–38^. Notably, ketamine can rapidly restore synaptic structure^81^. These morphological changes and subsequent functional deficits suggest that stress weakens excitatory synapses, and glutamatergic-modulating drugs, such as ketamine, may be beneficial in restoring synaptic strength.

Inflammation has been proposed as an indirect pathway connecting stress and glutamate^3, 70, 82^. PTSD is associated with increased pro-inflammatory markers and decreased anti-inflammatory markers^70, 82^. Pro-inflammatory cytokines may affect the glutamatergic system through the kynurenine pathway^70, 82^. Kynurenine breaks down into kynurenic acid in astrocytes and quinolinic acid in microglia. Kynurenic acid is an NMDA receptor antagonist that inhibits glutamate release, and quinolinic acid is an NMDA receptor agonist that activates glutamate release^70^. The kynurenine pathway has been implicated in other psychiatric disorders, including schizophrenia and mood disorders^83^. Glial pathology and inflammation^83^ may be related to the glutamate reductions that we observed in individuals with PTSD.

We observed lower NAA in the PTSD group compared to the group without trauma exposure. NAA is synthesized in neuronal mitochondria and is a putative marker of neuronal health and mitochondrial metabolism^84, 85^. NAA is linked to glutamate through the tricarboxylic acid and glutamate-glutamine cycles^84^ and is a precursor of NAAG, which regulates the release of glutamate, GABA, and other neurotransmitters^84,85^. The most consistent finding across PTSD MRS studies is reduced NAA in the ACC and bilateral hippocampi^43^. Studies of the DLPFC have found no difference in NAA between individuals with PTSD and trauma-exposed controls^41^ and no change following antidepressant treatment in individuals with comorbid PTSD and major depression^40^. Our finding of reduced NAA in individuals with PTSD is consistent with the volume reductions observed with structural neuroimaging in humans^86^ and the stress-induced morphological changes in the rodent PFC^73–79^.

We found elevated lactate in the PTSD group compared to the group without trauma exposure, which is consistent with PTSD metabolomics studies^87–89^. Similarly, studies of other psychiatric disorders have shown elevated lactate in individuals with depression^90^, schizophrenia^91, 92^, and bipolar disorder^93–97^ as well as individuals with panic disorder in response to hyperventilation^98^, lactate infusion^99, 100^, and visual stimulation^101, 102^. Elevated lactate is indicative of enhanced anaerobic glycolysis and may reflect impaired mitochondrial function, oxidative stress, or inflammation^87, 91, 93, 103,104^. Since the lactate MRS signal is small and overlaps with lipid signals, additional studies using an acquisition method optimized for lactate are needed to replicate our findings^104^.

A strength of our study is our use of 7T MRS, which allowed us to measure fourteen metabolites in a single acquisition. Unlike at lower field strengths where Glx (glutamate + glutamine) is the common measure, 7T’s higher spectral resolution enabled us to distinguish between glutamate and glutamine, and our finding was specific to glutamate. Similarly, we observed a difference in NAA but not tNAA (NAA + NAAG), which is the common measure at lower field strengths. Additionally, the higher signal-to-noise ratio at 7T allowed us to quantify smaller signals, such as GABA and glutathione. Detecting these metabolites at lower field strengths typically requires additional sequences and scan time. The high sensitivity of our findings highlights the advantage of 7T MRS compared to previous studies at lower field strengths.

Our findings should be considered in the context of several limitations. First, this study used self-report measures of trauma exposure and PTSD symptoms. Studies have shown that PCL-5 total scores correlate with severity scores from the Clinician-Administered PTSD Scale for *DSM-5*^105, 106^, but future studies would benefit from including structured interviews. Second, we only evaluated Criterion A status for the index events, so it is possible that NT participants witnessed or learned about a Criterion A event. We think this is unlikely because NT participants’ self-identified worst events (i.e., index events) did not meet Criterion A. Furthermore, 18/26 of the NT participants did not directly experience or witness a potential Criterion A event (n=7 did not endorse any LEC-5 events, n=5 endorsed only their non-Criterion A index event, and an additional n=6 did not endorse any directly experienced or witnessed events, only indirectly experienced events). Nevertheless, future studies should evaluate Criterion A exposure for all endorsed events, not just the index event. Third, although we excluded participants with a history of serious psychiatric disorders and substance use disorders, we did not exclude participants with other comorbid diagnoses such as attention-deficit/hyperactivity disorder, which is associated with neurometabolic abnormalities^107^. Fourth, we did not exclude participants who were taking psychotropic medications or who reported substance use. However, our secondary analyses indicated that the results were similar when we accounted for these confounds. Fifth, we measured metabolites only in the DLPFC. These data were collected as a part of a larger study, so time constraints prevented us from acquiring spectra from another brain region to investigate regional specificity. Sixth, macromolecules contribute to the broad background signal in short-TE spectra, which could impact the results. We used the standard macromolecule basis set included with LCModel^49^, but future studies should use individually acquired macromolecule spectra or an acquisition method that minimizes macromolecular signals. Finally, an inherent limitation of proton MRS measurements of glutamate is that we cannot distinguish between the neurotransmitter and metabolic pools of glutamate. The field would greatly benefit from additional ^13^C- MRS studies^72^ to better understand the nature of glutamatergic dysfunction in individuals with PTSD.

## CONCLUSION

The results of our study add to the growing evidence of glutamatergic dysfunction in individuals with PTSD. We also provide evidence of altered levels of NAA and lactate, which is consistent with prior studies of PTSD and other psychiatric disorders. Further research is needed to unravel the precise mechanisms underlying glutamate alterations in individuals with PTSD and their implications for the development and treatment of the disorder. Interventions that enhance glutamatergic function and promote neuronal health are promising targets for drug development. High-field MRS offers insight into the neurometabolic abnormalities associated with PTSD and is a powerful tool to probe trauma- and stress-related neurotransmission and metabolism *in vivo*.

## ACKNOWLEDGMENTS

We thank the participants who took part in this study. We thank Steven Nichols and Clayton Ridner for assistance with scanning, Sarah Comer for assistance with data entry and scanning, Julie Rodiek for administrative assistance, Dr. Nouha Salibi for optimizing the MRS acquisition, Dr. Ron Beyers for hardware and technical assistance, and Dr. Laura Rowland, Dr. Tom Denney, and Dr. Jeff Katz for helpful discussions. Data were presented at the annual meetings of the Society of Biological Psychiatry, the American College of Neuropsychopharmacology, and the International Society for Traumatic Stress Studies.

## AUTHOR CONTRIBUTIONS

Study concept and design: MAR. Acquisition, analysis, or interpretation of data: all authors. Statistical analysis: MAR. Drafting of the manuscript: MAR. Critical revision of the manuscript for important intellectual content: all authors. Administrative, technical, or material support: MAR, SEW, AAC, SMJ. Obtaining funding: MAR. Study supervision: MAR.

## STATEMENTS AND DECLARATIONS

*Ethical Considerations*

The Auburn University Institutional Review Board approved this study.

*Consent to Participate*

All participants provided written informed consent.

*Consent for Publication*

Not applicable

*Declaration of Conflicting Interests*

The authors declared no potential conflicts of interest with respect to the research, authorship, and/or publication of this article.

