## Supplementary Material for "Prefrontal metabolite alterations in individuals with posttraumatic stress disorder: a 7T magnetic resonance spectroscopy study"

for

**METHODS**

*Participants*

Fourteen additional people were enrolled in the study but excluded for claustrophobia (n = 3), dizziness (n = 1), image artifacts caused by dental work (n = 1), unclear *DSM-5* Criterion A status (n = 2), and psychiatric comorbidities (bipolar disorder: n = 3; obsessive-compulsive disorder: n = 2; antisocial personality disorder: n = 1; alcohol use disorder: n = 1).

Participants completed the Life Events Checklist for *DSM-5* (LEC-5) extended version^1^ to determine *DSM-5* Criterion A status. The LEC-5 is a self-report measure of trauma exposure with 17 categories of traumatic events. For each category, participants indicated how they experienced the event by selecting “happened to me,” “witnessed it,” “learned about it,” “part of my job,” “not sure,” or “doesn’t apply.” Participants were asked to select the event that they considered the worst overall and to describe the event. They were asked additional questions about this index event, including how long ago the event occurred, if someone’s life was in danger, if someone was seriously injured or killed, or if the event involved sexual violence. In addition to the LEC-5 questions, participants were asked if someone was threatened with serious physical harm and how old they were when the event happened. Index events were coded as meeting *DSM-5* Criterion A if participants:

(1) endorsed the event as “having happened to me directly” or “witnessed it”

AND

endorsed “my life was in danger,” “someone else’s life was in danger,” “I was seriously injured”, “someone else was seriously injured or killed,” “sexual violence,” “death of a close family member or close friend due to accident or violence,” “I was threatened with serious physical harm,” “someone else was threatened with serious physical harm,” or the written trauma narrative indicated a lack of consent in the case of sexual assault; or

(2) endorsed “learned about it happening to a close family member or close friend”

AND

endorsed “someone else was seriously injured or killed,” “sexual violence,” or “death of a close family member or close friend due to accident or violence”; or

(3) endorsed “part of my job”

AND

endorsed “someone else was seriously injured or killed.”

Participants’ written trauma narratives were used to verify Criterion A status. Two participants in the no trauma exposure (NT) group endorsed directly experiencing a transportation accident in addition to their non-Criterion A index events, but they provided sufficient details to rule out Criterion A trauma exposure.

The PTSD Checklist for *DSM-5­* (PCL-5)^1^ is a 20-item self-report measure to assess PTSD symptoms. Participants completed the PCL-5 based on their self-identified worst event (i.e., index event) by rating how much they were bothered by each symptom in the past month using the following scale: 0 = “not at all,” 1 = “a little bit,” 2 = “moderately,” 3 = “quite a bit,” and 4 = “extremely.” The PCL-5 total score is the sum of the item scores (range 0-80). PCL-5 items were dichotomized such that items rated as 2 (“moderately”) or higher were considered endorsed symptoms. A provisional *DSM-5* PTSD diagnosis can be made when participants endorse at least 1 re-experiencing symptom, 1 avoidance symptom, 2 symptoms of negative alterations in cognition and mood, and 2 hyperarousal symptoms. PCL-5 total scores of 31-33 and above have been shown to indicate probable PTSD, although studies have used cutoff scores ranging from 23 to 49, and a universal cutoff score has not been established^2^.

Information about alcohol, nicotine or tobacco, and other substance use was obtained for n = 64 participants using the *DSM-5* Self-Rated Level 1 Cross-Cutting Symptom Measure^3^, which is a 23-item self-report measure of 13 domains across psychiatric disorders. Participants rated how much or how often they were bothered by each problem in the past 2 weeks using the following scale: 0 = “none / not at all,” 1 = “slight / rare, less than a day or two,” 2 = “mild / several days,” 3 = “moderate / more than half the days,” 4 = “severe / nearly every day.” This measure was used only to determine current substance use and was not used to include/exclude participants, except for the secondary analysis in which all substance users were excluded (see Supplementary Results below).

*MR acquisition and analysis*

Data were acquired on a Siemens 7T MAGNETOM scanner equipped with a single-channel transmit and 32-channel receive head coil (Nova Medical) (Table S1). Images and spectra were acquired with the Siemens AutoAlign feature. Three-dimensional structural images were acquired for anatomical reference and MRS voxel segmentation (MPRAGE, TR/TE/TI = 2200/2.82/1050 ms, flip angle = 7°, field of view = 256 × 256 mm, matrix = 256 × 256, slice thickness = 1 mm, gap = 0.5 mm, 192 slices, GRAPPA acceleration factor = 2, sagittal acquisition).

Following FASTESTMAP shimming and voxel-based flip angle calibration^4^, spectra were acquired using an ultra-short TE STEAM sequence (TE/TR/TM = 5/10,000/45 ms, 32 averages, 4 kHz spectral bandwidth, 2048 points) with outer volume suppression and VAPOR water suppression^5-8^ (Table S1). The long TR was needed because of the increased power deposition resulting from the ultra-short TE, VAPOR water suppression, and outer volume suppression. A long TR and a short TE minimize the signal attenuation due to T1 and T2 effects, respectively. Spectra without water suppression (4 averages) were acquired for pre-processing and quantification. Spectra were coil-combined and averaged before being exported for processing. We previously demonstrated excellent reproducibility for glutamate with this approach^5^.

Spectra were processed with Osprey^9^ (version 2.4.0) and LCModel^10^ (version 6.3-1N implemented in Osprey; Table S1). Osprey pre-processing included the removal of the residual water signal and eddy current correction. Spectra were fit in LCModel in the range of 0.2-4.2 ppm. The simulated basis set^5^ included alanine, ascorbate, aspartate, creatine (Cr), gamma-aminobutyric acid (GABA), glucose, glutamine (Gln), glutamate (Glu), glycine, glycerophosphocholine (GPC), glutathione (GSH), lactate (Lac), myo-inositol (mI), *N*-acetylaspartate (NAA), *N*-acetylaspartylglutamate (NAAG), phosphocholine (PCh), phosphocreatine (PCr), phosphoethanolamine, scyllo-inositol, serine, and taurine. In Osprey, the knot spacing parameter bLineKnotSpace was set at 0.15, which is the default value for the LCModel parameter DKNTMN^11^. The LCModel parameters NSIMUL, CHCOMB, and CHOMIT were modified in the Osprey code to use the LCModel default settings. Glycerophosphocholine (GPC) and phosphocholine (PCh) were not reliably separated, so only the sum (tCho) is reported^5^.

*Statistical analysis*

Statistical analyses were conducted in RStudio version 2023.03.0+386 with R version 4.2.3 (2023-03-15)^12^. The following R packages were used: *emmeans* (version 1.8.5; <https://CRAN.R-project.org/package=emmeans>), *ggrain* (version 0.0.3; <https://CRAN.R-project.org/package=ggrain>)^13^, *here* (version 1.0.1; <https://CRAN.R-project.org/package=here>), *psych* (version 2.3.3; <https://CRAN.R-project.org/package=psych>), *rstatix* (version 0.7.2; <https://CRAN.R-project.org/package=rstatix>), *scales* (version 1.2.1; <https://CRAN.R-project.org/package=scales>), *stringr* (version 1.5.0; <https://CRAN.R-project.org/package=stringr>), *tableone* (version 0.13.2; <https://CRAN.R-project.org/package=tableone>), and *tidyverse* (version 2.0.0; <https://CRAN.R-project.org/package=tidyverse>)^14^.

**RESULTS**

We assessed the robustness of the glutamate, NAA, and lactate results after accounting for potential confounding factors. First, we examined the results after adding a covariate for psychotropic medication status (currently medicated or unmedicated). Medication status was not significant for glutamate (F_1,70_ = 0.50, p = 0.48, η^2^_p_ = 0.01), NAA (F_1,71_ = 0.02, p = 0.88, η^2^_p_ = 0.0003), and lactate (F_1,67_ = 1.76, p = 0.19, η^2^_p_ = 0.03), and the results did not meaningfully change (Table S9). Second, we examined the results after excluding TE and NT participants who endorsed more than two PTSD symptoms on the PCL-5 (i.e., more than two items rated “moderately” or higher) or had BDI or BAI scores higher than the “mild” range (BDI > 19 or BAI > 15). In other words, we retained only the minimally symptomatic TE participants. The results did not meaningfully change (Table S10). Third, we examined the results after excluding participants who reported using alcohol, nicotine or tobacco, or other substances (i.e., endorsed “slight / rare, less than a day or two” or higher on the substance use items of the *DSM-5* Self-Rated Level 1 Cross-Cutting Symptom Measure). Substance use information was not available for 16 participants, so these participants were also excluded from this secondary analysis. The difference in glutamate for PTSD compared to NT remained statistically significant, but the difference between TE and NT was no longer significant (Table S11). The difference in NAA between PTSD and NT remained statistically significant, and the difference between TE and NT became significant (Table S11). The overall group difference in lactate was no longer significant (Table S11). Fourth, we examined the results after including only the PTSD and TE participants who directly experienced or witnessed their index events (i.e., excluding “learned about” and “job” exposures). The results did not meaningfully change (Table S12). Fifth, we examined the results after reclassifying 4 TE participants as PTSD: 3 TE participants self-reported a PTSD diagnosis but did not meet *DSM-5* diagnostic criteria based on the PCL-5, and 1 TE participant met *DSM-5* diagnostic criteria but had a PCL-5 total score of less than 30. The results did not meaningfully change (Table S13). Finally, we examined the results after excluding 2 participants (1 TE and 1 NT) who had incidental findings on their structural MRI scans (near the left temporal lobe and the cerebellum) and 2 participants (1 PTSD and 1 TE) who reported unspecified movement disorders other than Parkinson’s or Huntington’s. The results did not meaningfully change (Table S14).

**Table S1.** Minimum reporting standards for *in vivo* magnetic resonance spectroscopy (MRSinMRS^15^)

| **MRSinMRS** | |
| --- | --- |
| **Site:** Auburn University MRI Research Center | |
| 1. **Hardware** | |
| a. Field strength | 7T |
| b. Manufacturer | Siemens |
| c. Model | MAGNETOM (software version VB17) |
| d. RF coil | ^1^H, single-channel transmit and 32-channel receive head coil (Nova Medical) |
| e. Additional hardware | N/A |
| 2. **Acquisition** | |
| a. Pulse sequence | STEAM (Siemens WIP 643) |
| b. Volume of interest (VOI) | left dorsolateral prefrontal cortex (Figure 1) |
| c. Nominal VOI size | 25 × 25 × 25 mm^3^ |
| d. Repetition time (TR), echo time (TE) | TR: 10,000 ms, TE: 5 ms |
| e. Total number of excitations or acquisitions per spectrum | 32 |
| f. Additional sequence parameters | TM: 45 ms, spectral bandwidth: 4000 Hz, number of points: 2048 |
| g. Water suppression method | VAPOR |
| h. Shimming method | FASTESTMAP followed by manual shimming of water |
| i. Triggering or motion correction method | N/A |
| 3. **Data analysis methods and outputs** | |
| a. Analysis software | Osprey (version 2.4.0), LCModel (version 6.3-1N implemented in Osprey), SPM12 |
| b. Processing steps deviating from quoted reference or product | Spectra were coil combined and averaged by the scanner software before being exported for processing. |
| c. Output measure | absolute concentration (molal: mmol/kg of tissue water; TissCorrWaterScaled as reported by Osprey) |
| d. Quantification references and assumptions, fitting model assumptions | Fit range: 0.2-4.2 ppm  Osprey bLineKnotSpace (LCModel DKNTMN): 0.15  LCModel parameters NSIMUL, CHCOMB, and CHOMIT were modified in the Osprey code to use the LCModel default settings.  Simulated basis set: alanine, ascorbate, aspartate, creatine (Cr), gamma-aminobutyric acid (GABA), glucose, glutamine (Gln), glutamate (Glu), glycine, glycerophosphocholine (GPC), glutathione (GSH), lactate (Lac), myo-inositol (mI), *N*-acetylaspartate (NAA), *N*-acetylaspartylglutamate (NAAG), phosphocholine (PCh), phosphocreatine (PCr), phosphoethanolamine, scyllo-inositol, serine, taurine, and LCModel’s default macromolecules |
| 4. **Data quality** | |
| a. Reported variables | creatine SNR, creatine linewidth, and water linewidth as reported by Osprey (Table 2) |
| b. Data exclusion criteria | poor fit, poor water suppression, water linewidth > 30 Hz, creatine linewidth > 30 Hz, CRLB > 20% (> 30% for lactate and NAAG) |
| c. Quality measures of postprocessing model fitting | Cramer-Rao lower bounds as reported by LCModel (Table 2) |
| d. Sample spectrum | Figure 1 and Figure S1 |

**Table S2.** Number of participants excluded from statistical analyses

|  | **PTSD** | **TE** | **NT** |
| --- | --- | --- | --- |
| Sample size | 27 | 27 | 26 |
| Participants excluded for poor spectral quality | 1 | 0 | 2 |
| Sample size entered into statistical analyses | 26 | 27 | 24 |
| Participants excluded based on CRLB threshold |  |  |  |
| GABA | 1 | 6 | 2 |
| NAAG | 0 | 0 | 1 |
| PCr | 2 | 1 | 1 |
| Lac | 1 | 3 | 0 |
| Participants excluded as statistical outliers |  |  |  |
| Glu | 0 | 1 | 0 |
| Gln | 1 | 0 | 0 |
| Glx | 0 | 1 | 0 |
| GABA | 1 | 0 | 0 |
| NAAG | 0 | 1 | 0 |
| GSH | 0 | 1 | 0 |

Abbreviations: CRLB, Cramer-Rao lower bounds; GABA, gamma-aminobutyric acid; Gln, glutamine; Glu, glutamate; Glx, glutamate + glutamine; GSH, glutathione; Lac, lactate; NAAG, N-acetylaspartylglutamate; NT, no trauma exposure; PCr, phosphocreatine; PTSD, posttraumatic stress disorder; TE, trauma-exposed without PTSD

**Table S3.** Post hoc pairwise comparisons of clinical characteristics

|  | **t-statistic** | **p-value** |
| --- | --- | --- |
| BDI |  |  |
| PTSD vs. TE | 8.79 | p_Tukey_ < 0.001 |
| PTSD vs. NT | 10.20 | p_Tukey_ < 0.001 |
| TE vs. NT | 1.49 | p_Tukey_ = 0.30 |
| BAI |  |  |
| PTSD vs. TE | 8.75 | p_Tukey_ < 0.001 |
| PTSD vs. NT | 11.42 | p_Tukey_ < 0.001 |
| TE vs. NT | 2.76 | p_Tukey_ = 0.02 |
| MDI |  |  |
| PTSD vs. TE | 8.16 | p_Tukey_ < 0.001 |
| PTSD vs. NT | 9.73 | p_Tukey_ < 0.001 |
| TE vs. NT | 1.64 | p_Tukey_ = 0.23 |
| Comorbid disorders |  |  |
| PTSD vs. TE | --- | p_Bonferroni_ = 0.26 |
| PTSD vs. NT | --- | p_Bonferroni_ < 0.001 |
| TE vs. NT | --- | p_Bonferroni_ = 0.18 |
| Psychotropic medications |  |  |
| PTSD vs. TE | --- | p_Bonferroni_ = 0.41 |
| PTSD vs. NT | --- | p_Bonferroni_ = 0.04 |
| TE vs. NT | --- | p_Bonferroni_ > 0.99 |

Abbreviations: BAI, Beck Anxiety Inventory; BDI, Beck Depression Inventory; MDI, Multiscale Dissociation Inventory; NT, no trauma exposure; PTSD, posttraumatic stress disorder; TE, trauma-exposed without PTSD

**Table S4.** Post hoc pairwise comparisons of tissue fractions

|  | **t-statistic** | **p_Tukey_** |
| --- | --- | --- |
| Gray matter |  |  |
| PTSD vs. TE | 0.04 | > 0.99 |
| PTSD vs. NT | 2.34 | 0.06 |
| TE vs. NT | 2.40 | 0.049 |
| White matter |  |  |
| PTSD vs. TE | 0.02 | > 0.99 |
| PTSD vs. NT | 2.40 | 0.049 |
| TE vs. NT | 2.44 | 0.04 |

Abbreviations: NT, no trauma exposure; PTSD, posttraumatic stress disorder; TE, trauma-exposed without PTSD

**Table S5.** Post hoc pairwise comparisons of metabolite concentrations

|  | **t-statistic** | **p_Tukey_** | **Cohen’s d** |
| --- | --- | --- | --- |
| Glu |  |  |  |
| PTSD vs. TE | 0.43 | 0.90 | 0.13 |
| PTSD vs. NT | 3.23 | 0.005 | 0.92 |
| TE vs. NT | 2.78 | 0.02 | 0.80 |
| NAA |  |  |  |
| PTSD vs. TE | 1.29 | 0.41 | 0.37 |
| PTSD vs. NT | 3.47 | 0.002 | 0.99 |
| TE vs. NT | 2.21 | 0.08 | 0.63 |
| Lac |  |  |  |
| PTSD vs. TE | 1.90 | 0.15 | 0.56 |
| PTSD vs. NT | 3.51 | 0.002 | 1.01 |
| TE vs. NT | 1.53 | 0.28 | 0.45 |

Abbreviations: Glu, glutamate; Lac, lactate; NAA, *N*-acetylaspartate; NT, no trauma exposure; PTSD, posttraumatic stress disorder; TE, trauma-exposed without PTSD

**Table S6.** Post hoc pairwise comparisons of metabolite concentrations after alpha correction

|  | **t-statistic** | **p_Tukey_** | **Cohen’s d** |
| --- | --- | --- | --- |
| Glu |  |  |  |
| PTSD vs. TE | 0.44 | 0.90 | 0.13 |
| PTSD vs. NT | 3.23 | 0.005 | 0.93 |
| TE vs. NT | 2.78 | 0.02 | 0.80 |

Abbreviations: Glu, glutamate; NT, no trauma exposure; PTSD, posttraumatic stress disorder; TE, trauma-exposed without PTSD

**Table S7.** Post hoc pairwise comparisons of metabolite concentrations after including tissue fraction as a covariate

|  | **t-statistic** | **p_Tukey_** | **Cohen’s d** |
| --- | --- | --- | --- |
| Glu |  |  |  |
| PTSD vs. TE | 0.15 | 0.99 | 0.05 |
| PTSD vs. NT | 2.10 | 0.10 | 0.65 |
| TE vs. NT | 2.02 | 0.12 | 0.60 |
| NAA |  |  |  |
| PTSD vs. TE | 0.94 | 0.62 | 0.27 |
| PTSD vs. NT | 2.41 | 0.048 | 0.74 |
| TE vs. NT | 1.62 | 0.24 | 0.47 |
| Lac |  |  |  |
| PTSD vs. TE | 1.80 | 0.18 | 0.54 |
| PTSD vs. NT | 2.89 | 0.01 | 0.90 |
| TE vs. NT | 1.18 | 0.47 | 0.36 |

Abbreviations: Glu, glutamate; Lac, lactate; NAA, *N*-acetylaspartate; NT, no trauma exposure; PTSD, posttraumatic stress disorder; TE, trauma-exposed without PTSD

**Table S8.** Metabolite comparisons between groups whose index trauma occurred in adulthood vs. childhood/adolescence (combined sample of PTSD and TE participants)

|  | **Adulthood** | **Childhood/Adolescence** | **Statistic** | **p-value** | **η^2^_p_** |
| --- | --- | --- | --- | --- | --- |
| Glu | 9.85 (0.67) | 10.02 (0.72) | F_1,48_ = 0.30 | 0.59 | 0.006 |
| n | 26 | 26 |  |  |  |
| CRLB, % | 2.00 (0.00) | 2.04 (0.20) | F_1,50_ = 1.00 | 0.35 |  |
| NAA | 12.60 (0.69) | 12.81 (1.02) | F_1,49_ = 0.00004 | > 0.99 | 0.000001 |
| n | 26 | 27 |  |  |  |
| CRLB, % | 2.04 (0.20) | 2.00 (0.39) | F_1,51_ = 0.20 | 0.66 |  |
| Lac | 1.34 (0.34) | 1.20 (0.32) | F_1,45_ = 0.64 | 0.43 | 0.01 |
| n | 25 | 24 |  |  |  |
| CRLB, % | 15.00 (3.28) | 15.04 (3.72) | F_1,47_ = 0.002 | 0.97 |  |

Abbreviations: CRLB, Cramer-Rao lower bounds; Glu, glutamate; Lac, lactate; NAA, *N*-acetylaspartate; PTSD, posttraumatic stress disorder; TE, trauma-exposed without PTSD

**Table S9.** Metabolite comparisons with an additional covariate for psychotropic medication status

|  | **PTSD** | **TE** | **NT** | **Statistic** | **p-value** | **Effect size** |
| --- | --- | --- | --- | --- | --- | --- |
| Glu | 9.95 (0.70) | 9.91 (0.70) | 10.51 (0.72) | F_2,70_ = 5.07 | p = 0.009 | η^2^_p_ = 0.13 |
| n | 26 | 26 | 24 |  |  |  |
| PTSD vs. TE |  |  |  | t = 0.26 | p_Tukey_ = 0.96 | d = 0.08 |
| PTSD vs. NT |  |  |  | t = 2.81 | p_Tukey_ = 0.02 | d = 0.85 |
| TE vs. NT |  |  |  | t = 2.68 | p_Tukey_ = 0.02 | d = 0.77 |
| NAA | 12.61 (0.81) | 12.80 (0.93) | 13.36 (0.69) | F_2,71_ = 5.68 | p = 0.005 | η^2^_p_ = 0.14 |
| n | 26 | 27 | 24 |  |  |  |
| PTSD vs. TE |  |  |  | t = 1.27 | p_Tukey_ = 0.41 | d = 0.38 |
| PTSD vs. NT |  |  |  | t = 3.31 | p_Tukey_ = 0.004 | d = 1.00 |
| TE vs. NT |  |  |  | t = 2.20 | p_Tukey_ = 0.08 | d = 0.63 |
| Lac | 1.34 (0.35) | 1.20 (0.31) | 1.03 (0.28) | F_2,67_ = 4.33 | p = 0.02 | η^2^_p_ = 0.11 |
| n | 25 | 24 | 24 |  |  |  |
| PTSD vs. TE |  |  |  | t = 1.60 | p_Tukey_ = 0.25 | d = 0.48 |
| PTSD vs. NT |  |  |  | t = 2.94 | p_Tukey_ = 0.01 | d = 0.89 |
| TE vs. NT |  |  |  | t = 1.39 | p_Tukey_ = 0.35 | d = 0.41 |

Abbreviations: Glu, glutamate; Lac, lactate; NAA, *N*-acetylaspartate; NT, no trauma exposure; PTSD, posttraumatic stress disorder; TE, trauma-exposed without PTSD

**Table S10.** Metabolite comparisons after excluding TE and NT participants who endorsed more than two PTSD symptoms or had greater than mild depression or anxiety symptoms

|  | **PTSD** | **TE** | **NT** | **Statistic** | **p-value** | **Effect size** |
| --- | --- | --- | --- | --- | --- | --- |
| Glu | 9.95 (0.70) | 9.92 (0.86) | 10.49 (0.73) | F_2,57_ = 5.59 | p = 0.006 | η^2^_p_ = 0.16 |
| n | 26 | 15 | 21 |  |  |  |
| PTSD vs. TE |  |  |  | t = 0.08 | p_Tukey_ > 0.99 | d = 0.03 |
| PTSD vs. NT |  |  |  | t = 2.99 | p_Tukey_ = 0.01 | d = 0.89 |
| TE vs. NT |  |  |  | t = 2.69 | p_Tukey_ = 0.03 | d = 0.92 |
| NAA | 12.61 (0.81) | 12.75 (0.72) | 13.30 (0.69) | F_2,57_ = 6.08 | p = 0.004 | η^2^_p_ = 0.18 |
| n | 26 | 15 | 21 |  |  |  |
| PTSD vs. TE |  |  |  | t = 0.42 | p_Tukey_ = 0.91 | d = 0.14 |
| PTSD vs. NT |  |  |  | t = 3.31 | p_Tukey_ = 0.005 | d = 0.98 |
| TE vs. NT |  |  |  | t = 2.47 | p_Tukey_ = 0.04 | d = 0.84 |
| Lac | 1.34 (0.35) | 1.23 (0.31) | 1.05 (0.30) | F_2,55_ = 5.05 | p = 0.010 | η^2^_p_ = 0.16 |
| n | 25 | 14 | 21 |  |  |  |
| PTSD vs. TE |  |  |  | t = 1.10 | p_Tukey_ = 0.52 | d = 0.38 |
| PTSD vs. NT |  |  |  | t = 3.17 | p_Tukey_ = 0.007 | d = 0.95 |
| TE vs. NT |  |  |  | t = 1.64 | p_Tukey_ = 0.24 | d = 0.57 |

Abbreviations: Glu, glutamate; Lac, lactate; NAA, *N*-acetylaspartate; NT, no trauma exposure; PTSD, posttraumatic stress disorder; TE, trauma-exposed without PTSD

**Table S11.** Metabolite comparisons after excluding participants who reported substance use^a^

|  | **PTSD** | **TE** | **NT** | **Statistic** | **p-value** | **Effect size** |
| --- | --- | --- | --- | --- | --- | --- |
| Glu | 9.88 (0.50) | 9.93 (0.73) | 10.54 (0.80) | F_2,40_ = 4.40 | p = 0.02 | η^2^_p_ = 0.18 |
| n | 15 | 17 | 13 |  |  |  |
| PTSD vs. TE |  |  |  | t = 0.96 | p_Tukey_ = 0.61 | d = 0.36 |
| PTSD vs. NT |  |  |  | t = 2.90 | p_Tukey_ = 0.02 | d = 1.12 |
| TE vs. NT |  |  |  | t = 2.02 | p_Tukey_ = 0.12 | d = 0.76 |
| NAA | 12.41 (0.76) | 12.62 (0.74) | 13.39 (0.76) | F_2,40_ = 6.46 | p = 0.004 | η^2^_p_ = 0.24 |
| n | 15 | 17 | 13 |  |  |  |
| PTSD vs. TE |  |  |  | t = 1.06 | p_Tukey_ = 0.54 | d = 0.40 |
| PTSD vs. NT |  |  |  | t = 3.50 | p_Tukey_ = 0.003 | d = 1.34 |
| TE vs. NT |  |  |  | t = 2.53 | p_Tukey_ = 0.04 | d = 0.95 |
| Lac | 1.37 (0.29) | 1.20 (0.31) | 1.18 (0.27) | F_2,39_ = 2.94 | p = 0.07 | η^2^_p_ = 0.13 |
| n | 15 | 16 | 13 |  |  |  |

Abbreviations: Glu, glutamate; Lac, lactate; NAA, *N*-acetylaspartate; NT, no trauma exposure; PTSD, posttraumatic stress disorder; TE, trauma-exposed without PTSD

^a^Substance use information was not available for n = 16 participants, so they were also excluded from this analysis.

**Table S12.** Metabolite comparisons when including only the PTSD and TE participants who directly experienced or witnessed traumatic events

|  | **PTSD** | **TE** | **NT** | **Statistic** | **p-value** | **Effect size** |
| --- | --- | --- | --- | --- | --- | --- |
| Glu | 9.91 (0.70) | 9.85 (0.64) | 10.51 (0.72) | F_2,58_ = 7.04 | p = 0.002 | η^2^_p_ = 0.20 |
| n | 21 | 18 | 24 |  |  |  |
| PTSD vs. TE |  |  |  | t = 0.22 | p_Tukey_ = 0.97 | d = 0.08 |
| PTSD vs. NT |  |  |  | t = 3.23 | p_Tukey_ = 0.004 | d = 1.01 |
| TE vs. NT |  |  |  | t = 2.89 | p_Tukey_ = 0.01 | d = 0.93 |
| NAA | 12.42 (0.75) | 12.87 (0.95) | 13.36 (0.69) | F_2,59_ = 9.27 | p < 0.001 | η^2^_p_ = 0.24 |
| n | 21 | 19 | 24 |  |  |  |
| PTSD vs. TE |  |  |  | t = 2.21 | p_Tukey_ = 0.08 | d = 0.74 |
| PTSD vs. NT |  |  |  | t = 4.31 | p_Tukey_ < 0.001 | d = 1.30 |
| TE vs. NT |  |  |  | t = 1.79 | p_Tukey_ = 0.18 | d = 0.56 |
| Lac | 1.32 (0.36) | 1.17 (0.30) | 1.03 (0.28) | F_2,55_ = 5.07 | p = 0.010 | η^2^_p_ = 0.16 |
| n | 20 | 16 | 24 |  |  |  |
| PTSD vs. TE |  |  |  | t = 1.78 | p_Tukey_ = 0.19 | d = 0.64 |
| PTSD vs. NT |  |  |  | t = 3.18 | p_Tukey_ = 0.007 | d = 0.97 |
| TE vs. NT |  |  |  | t = 1.02 | p_Tukey_ = 0.57 | d = 0.34 |

Abbreviations: Glu, glutamate; Lac, lactate; NAA, *N*-acetylaspartate; NT, no trauma exposure; PTSD, posttraumatic stress disorder; TE, trauma-exposed without PTSD

**Table S13.** Metabolite comparisons after reclassifying four TE participants as PTSD^a^

|  | **PTSD** | **TE** | **NT** | **Statistic** | **p-value** | **Effect size** |
| --- | --- | --- | --- | --- | --- | --- |
| Glu | 9.97 (0.65) | 9.88 (0.76) | 10.51 (0.72) | F_2,71_ = 6.09 | p = 0.004 | η^2^_p_ = 0.15 |
| n | 30 | 22 | 24 |  |  |  |
| PTSD vs. TE |  |  |  | t = 0.23 | p_Tukey_ = 0.97 | d = 0.07 |
| PTSD vs. NT |  |  |  | t = 3.01 | p_Tukey_ = 0.01 | d = 0.83 |
| TE vs. NT |  |  |  | t = 3.00 | p_Tukey_ = 0.01 | d = 0.90 |
| NAA | 12.61 (0.78) | 12.84 (0.98) | 13.36 (0.69) | F_2,72_ = 6.10 | p = 0.004 | η^2^_p_ = 0.15 |
| n | 30 | 23 | 24 |  |  |  |
| PTSD vs. TE |  |  |  | t = 1.22 | p_Tukey_ = 0.45 | d = 0.35 |
| PTSD vs. NT |  |  |  | t = 3.47 | p_Tukey_ = 0.003 | d = 0.96 |
| TE vs. NT |  |  |  | t = 2.07 | p_Tukey_ = 0.10 | d = 0.61 |
| Lac | 1.34 (0.33) | 1.16 (0.32) | 1.03 (0.28) | F_2,68_ = 6.72 | p = 0.002 | η^2^_p_ = 0.17 |
| n | 29 | 20 | 24 |  |  |  |
| PTSD vs. TE |  |  |  | t = 2.15 | p_Tukey_ = 0.09 | d = 0.64 |
| PTSD vs. NT |  |  |  | t = 3.62 | p_Tukey_ = 0.002 | d = 1.01 |
| TE vs. NT |  |  |  | t = 1.18 | p_Tukey_ = 0.47 | d = 0.36 |

Abbreviations: Glu, glutamate; Lac, lactate; NAA, *N*-acetylaspartate; NT, no trauma exposure; PTSD, posttraumatic stress disorder; TE, trauma-exposed without PTSD

^a^3 TE participants self-reported a PTSD diagnosis but did not meet *DSM-5* criteria based on the PCL-5, and 1 TE participant met *DSM-5* diagnostic criteria but had a PCL-5 total score of less than 30 (see inclusion criteria in the Methods).

**Table S14.** Metabolite comparisons after excluding participants with incidental findings and participants who reported unspecified movement disorders^a^

|  | **PTSD** | **TE** | **NT** | **Statistic** | **p-value** | **Effect size** |
| --- | --- | --- | --- | --- | --- | --- |
| Glu | 9.95 (0.71) | 9.91 (0.73) | 10.54 (0.72) | F_2,67_ = 6.36 | p = 0.003 | η^2^_p_ = 0.16 |
| n | 25 | 24 | 23 |  |  |  |
| PTSD vs. TE |  |  |  | t = 0.36 | p_Tukey_ = 0.93 | d = 0.11 |
| PTSD vs. NT |  |  |  | t = 3.26 | p_Tukey_ = 0.005 | d = 0.95 |
| TE vs. NT |  |  |  | t = 2.85 | p_Tukey_ = 0.02 | d = 0.84 |
| NAA | 12.67 (0.76) | 12.87 (0.93) | 13.41 (0.66) | F_2,68_ = 5.98 | p = 0.004 | η^2^_p_ = 0.15 |
| n | 25 | 25 | 23 |  |  |  |
| PTSD vs. TE |  |  |  | t = 1.20 | p_Tukey_ = 0.46 | d = 0.35 |
| PTSD vs. NT |  |  |  | t = 3.41 | p_Tukey_ = 0.003 | d = 0.99 |
| TE vs. NT |  |  |  | t = 2.21 | p_Tukey_ = 0.08 | d = 0.64 |
| Lac | 1.33 (0.35) | 1.19 (0.31) | 1.02 (0.28) | F_2,64_ = 6.21 | p = 0.003 | η^2^_p_ = 0.16 |
| n | 24 | 22 | 23 |  |  |  |
| PTSD vs. TE |  |  |  | t = 1.86 | p_Tukey_ = 0.16 | d = 0.56 |
| PTSD vs. NT |  |  |  | t = 3.52 | p_Tukey_ = 0.002 | d = 1.03 |
| TE vs. NT |  |  |  | t = 1.56 | p_Tukey_ = 0.27 | d = 0.47 |

Abbreviations: Glu, glutamate; Lac, lactate; NAA, *N*-acetylaspartate; NT, no trauma exposure; PTSD, posttraumatic stress disorder; TE, trauma-exposed without PTSD

^a^2 participants (1 TE and 1 NT) had incidental findings on their structural MRI scans (near the left temporal lobe and the cerebellum), and 2 participants (1 PTSD and 1 TE) reported unspecified movement disorders other than Parkinson’s or Huntington’s.

**Table S15.** Metabolite comparisons without the removal of outliers^a^

|  | **PTSD** | **TE** | **NT** | **Statistic** | **p-value** | **Effect size** |
| --- | --- | --- | --- | --- | --- | --- |
| Glu | 9.95 (0.70) | 10.02 (0.89) | 10.51 (0.72) | F_2,72_ = 4.85 | p = 0.01 | η^2^_p_ = 0.12 |
| n | 26 | 27 | 24 |  |  |  |
| CRLB, % | 2.04 (0.20) | 2.00 (0.00) | 2.13 (0.34) | F_2,74_ = 2.12 | p = 0.13 |  |
| PTSD vs. TE |  |  |  | t = 1.07 | p_Tukey_ = 0.54 | d = 0.30 |
| PTSD vs. NT |  |  |  | t = 3.06 | p_Tukey_ = 0.009 | d = 0.88 |
| TE vs. NT |  |  |  | t = 2.02 | p_Tukey_ = 0.11 | d = 0.57 |
| Gln | 2.31 (0.59) | 2.32 (0.40) | 2.33 (0.42) | F_2,72_ = 0.28 | p = 0.76 | η^2^_p_ = 0.01 |
| n | 26 | 27 | 24 |  |  |  |
| CRLB, % | 7.69 (2.13) | 7.56 (2.21) | 7.63 (2.14) | F_2,74_ = 0.03 | p = 0.97 |  |
| Glx | 12.18 (0.90) | 12.25 (1.10) | 12.74 (0.86) | F_2,72_ = 2.64 | p = 0.08 | η^2^_p_ = 0.07 |
| n | 26 | 27 | 24 |  |  |  |
| CRLB, % | 2.35 (0.49) | 2.48 (0.51) | 2.25 (0.53) | F_2,74_ = 1.34 | p = 0.27 |  |
| GABA | 1.50 (0.36) | 1.40 (0.27) | 1.50 (0.27) | F_2,63_ = 0.70 | p = 0.50 | η^2^_p_ = 0.02 |
| n | 25 | 21 | 22 |  |  |  |
| CRLB, % | 9.80 (2.47) | 10.43 (2.69) | 9.86 (2.27) | F_2,65_ = 0.43 | p = 0.65 |  |
| NAAG | 2.04 (0.46) | 2.07 (0.60) | 1.82 (0.57) | F_2,71_ = 1.46 | p = 0.24 | η^2^_p_ = 0.04 |
| n | 26 | 27 | 23 |  |  |  |
| CRLB, % | 8.96 (3.13) | 9.48 (3.82) | 11.09 (4.88) | F_2,73_ = 1.88 | p = 0.16 |  |
| GSH | 2.26 (0.31) | 2.20 (0.36) | 2.33 (0.35) | F_2,72_ = 1.64 | p = 0.20 | η^2^_p_ = 0.04 |
| n | 26 | 27 | 24 |  |  |  |
| CRLB, % | 4.92 (0.98) | 5.04 (0.98) | 4.96 (1.00) | F_2,74_ = 0.09 | p = 0.91 |  |

Abbreviations: CRLB, Cramer-Rao lower bounds; GABA, gamma-aminobutyric acid; Gln, glutamine; Glu, glutamate; Glx, glutamate + glutamine; GSH, glutathione; NAAG, *N*-acetylaspartylglutamate; NT, no trauma exposure; PTSD, posttraumatic stress disorder; TE, trauma-exposed without PTSD

^a^No outliers for NAA, tNAA, tCho, Cr, PCr, tCr, Lac, and mI (see outlier criterion in the Methods)
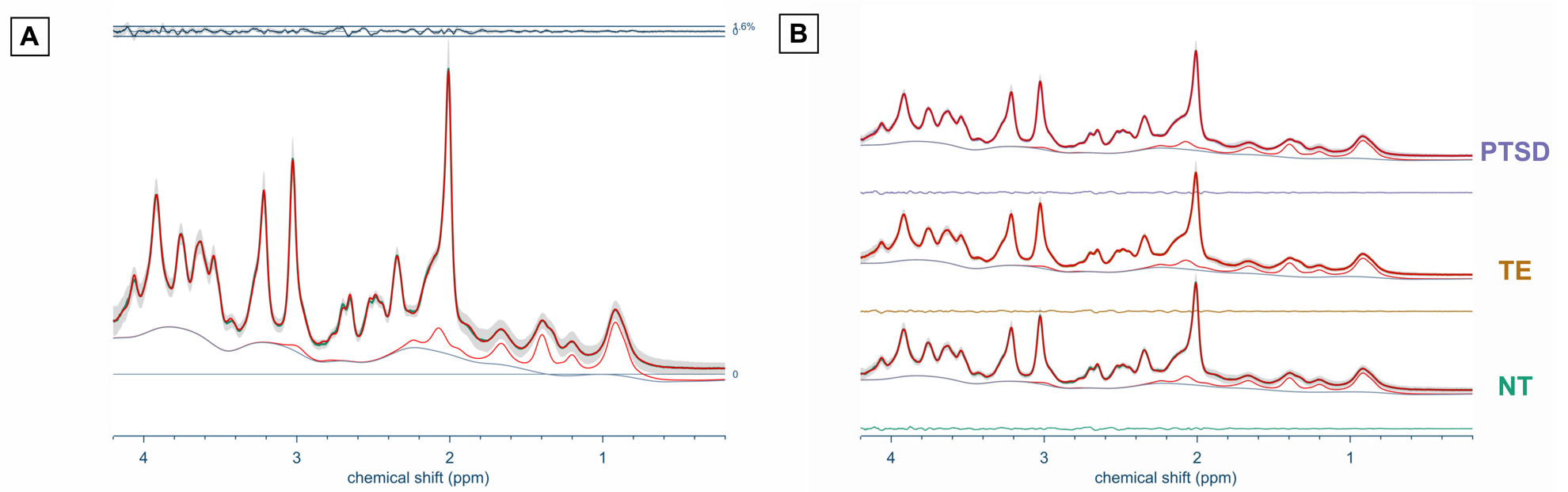

**Figure S1.** Group-averaged spectra in (A) the combined sample and (B) the samples of PTSD, trauma-exposed without PTSD (TE), and no trauma exposure (NT)

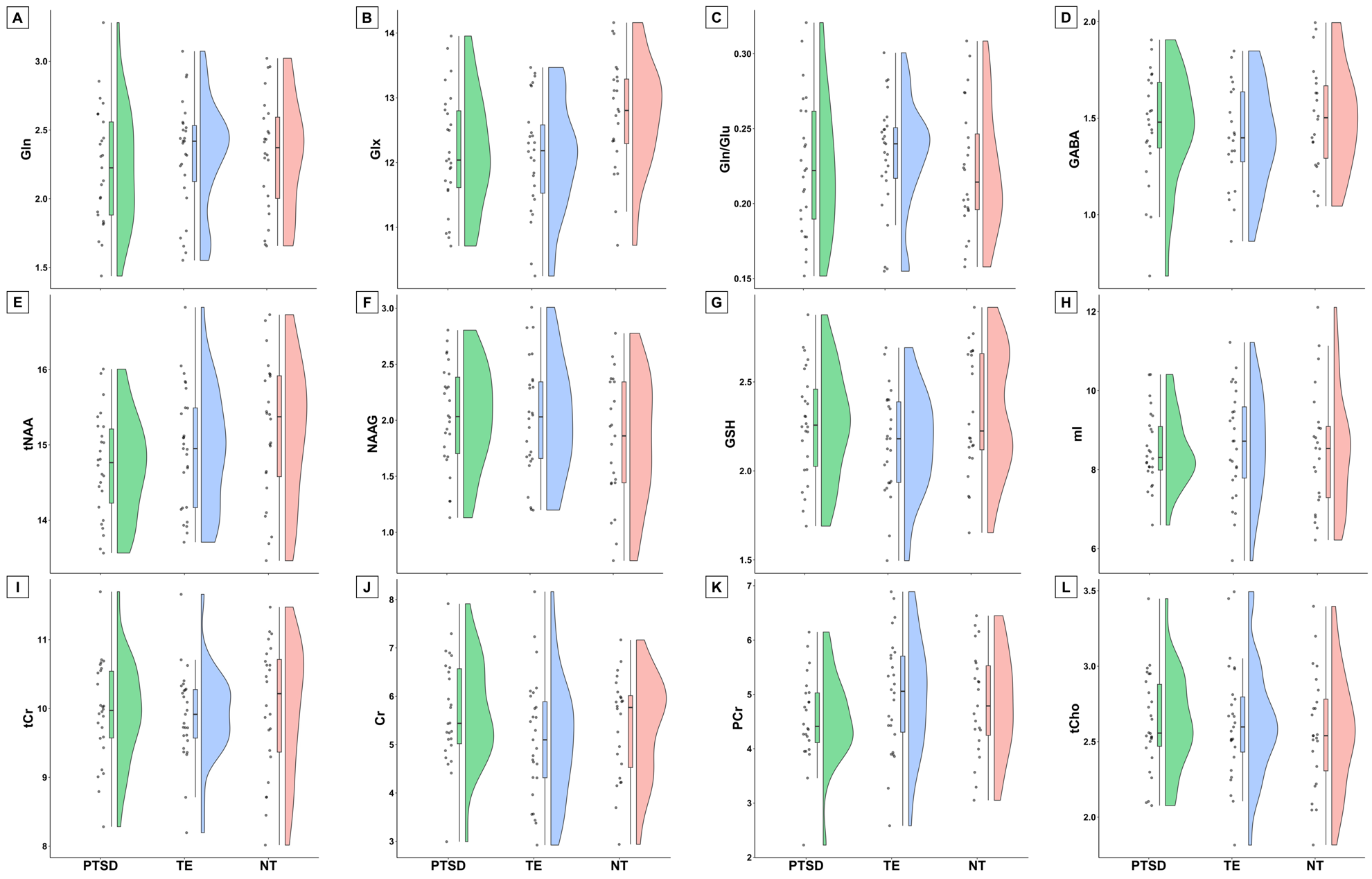

**Figure S2.** Additional metabolites (mmol/kg of tissue water): (A) glutamine, (B) Glx (glutamate + glutamine), (C) glutamine/glutamate ratio, (D) gamma-aminobutyric acid, (E) tNAA (*N*-acetylaspartate + *N*-acetylaspartylglutamate), (F) *N*-acetylaspartylglutamate, (G) glutathione, (H) myo-inositol (I) tCr (creatine + phosphocreatine), (J) creatine, (K) phosphocreatine, (L) tCho (glycerophosphocholine + phosphocholine)

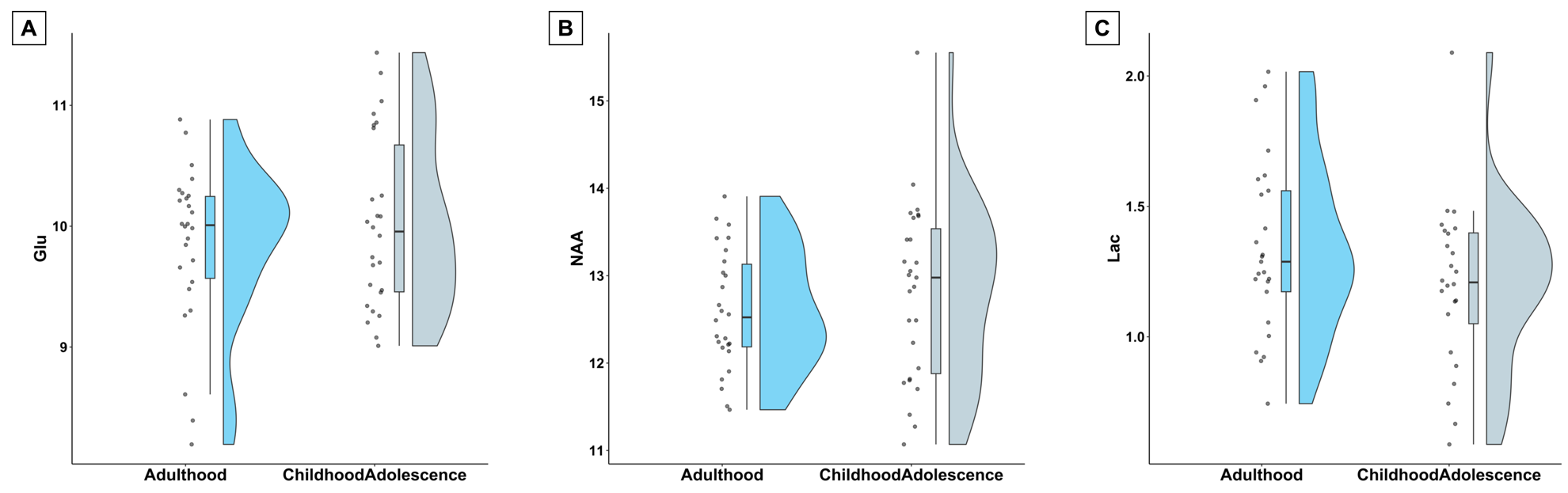

**Figure S3.** Comparison of (A) glutamate, (B) *N*-acetylaspartate, and (C) lactate concentrations (mmol/kg of tissue water) between groups whose index trauma occurred in adulthood vs. childhood/adolescence (combined sample of PTSD and trauma-exposed participants)

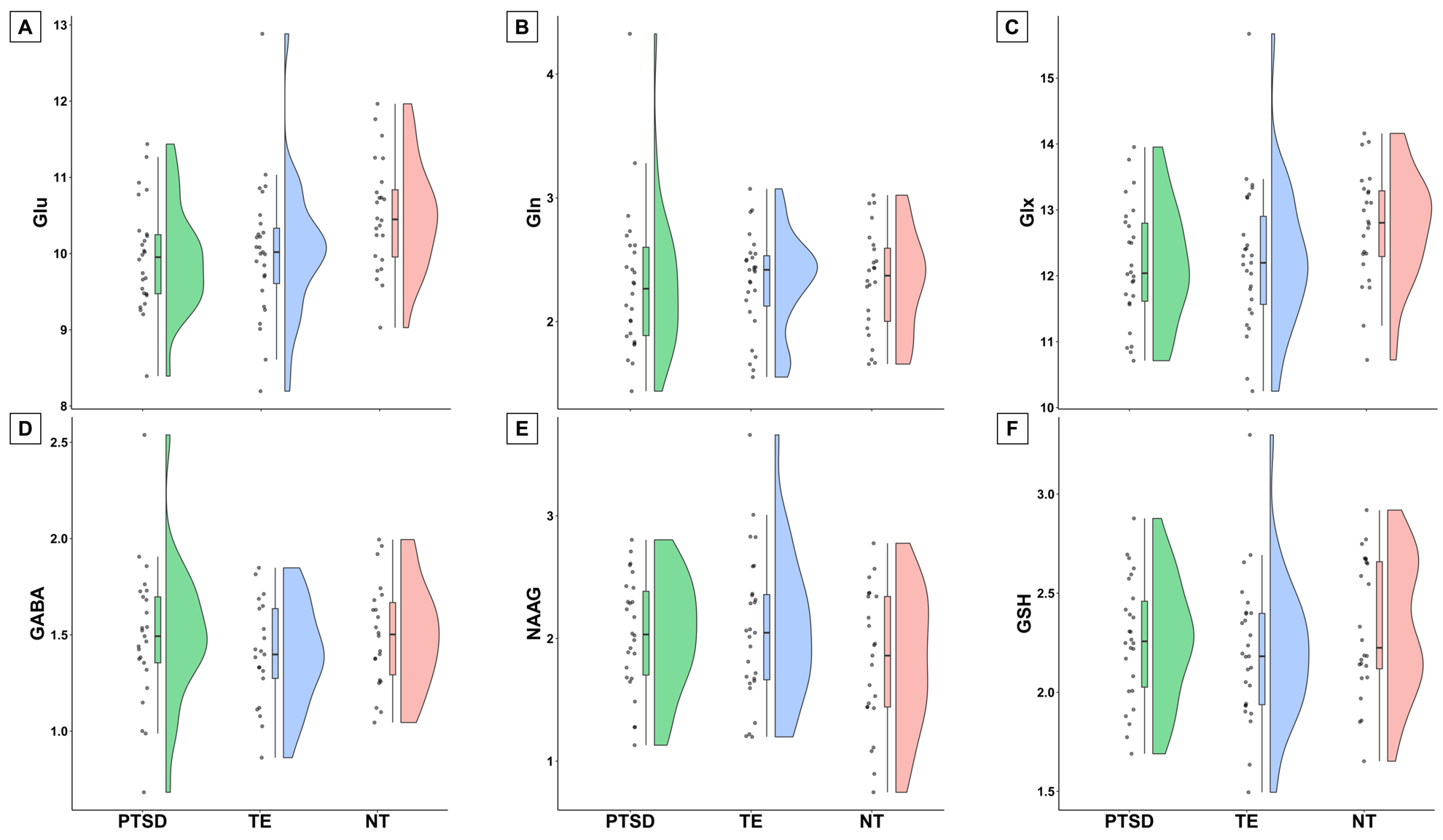

**Figure S4.** Metabolites without the removal of outliers (mmol/kg of tissue water): (A) glutamate, (B) glutamine, (C) Glx (glutamate + glutamine), (D) gamma-aminobutyric acid, (E) *N*-acetylaspartylglutamate, (F) glutathione. There were no outliers for NAA, tNAA, tCho, Cr, PCr, tCr, Lac, and mI.

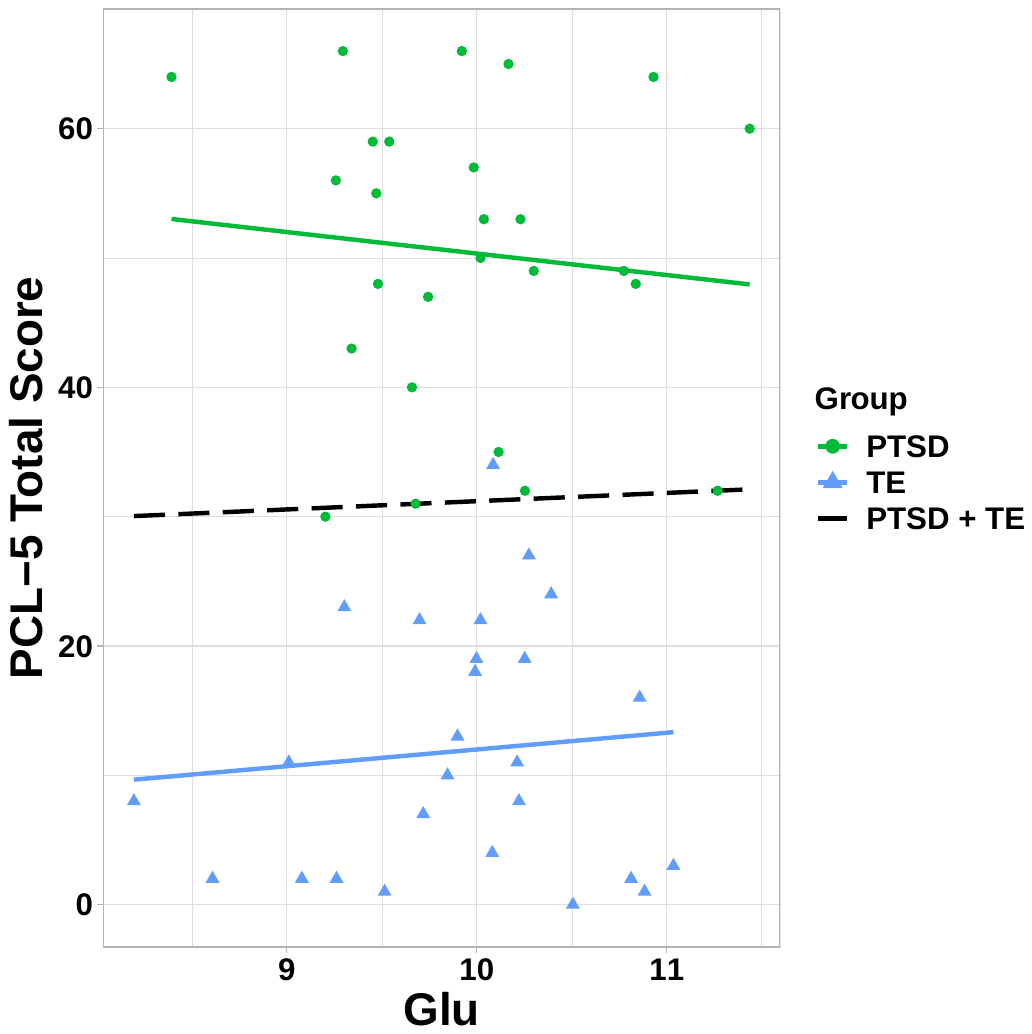

**Figure S5.** Correlation between glutamate (mmol/kg of tissue water) and PCL-5 total score
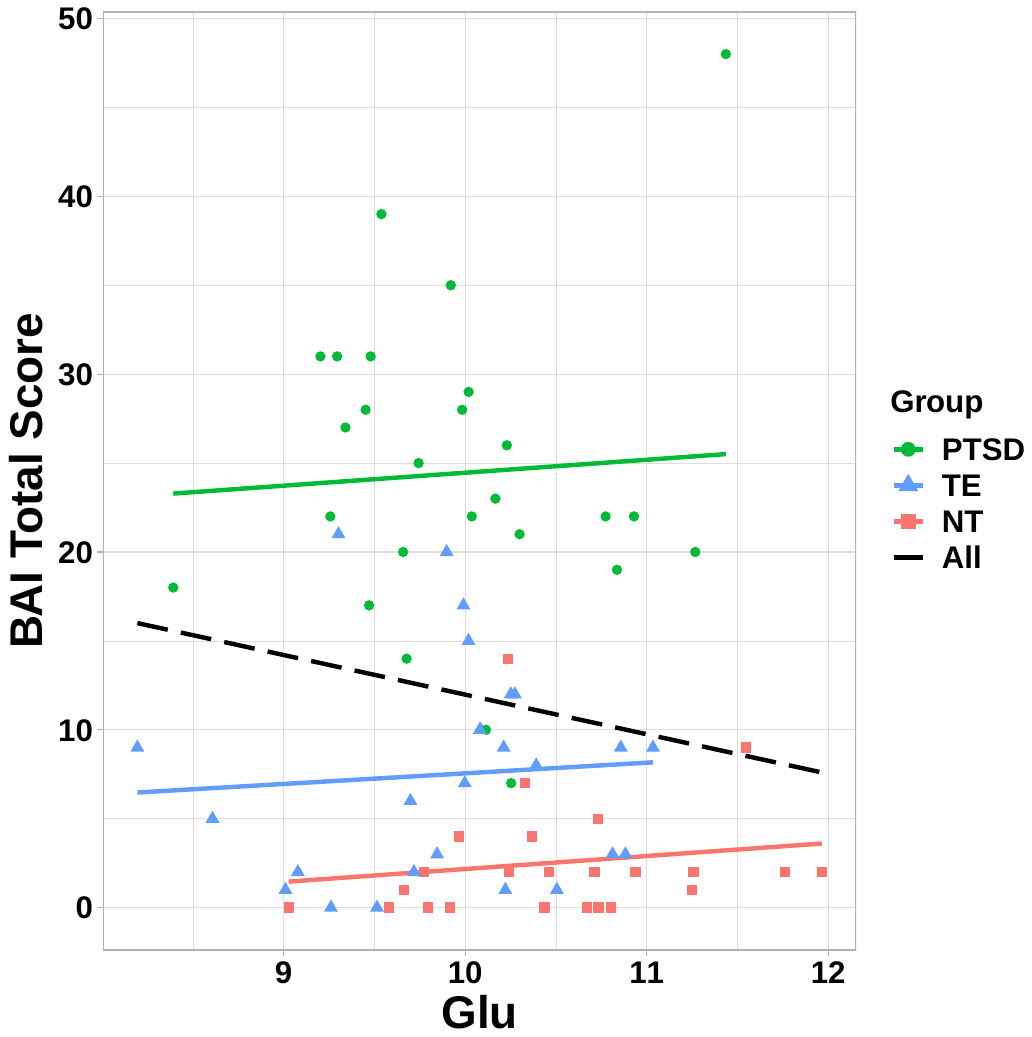

**Figure S6.** Correlation between glutamate (mmol/kg of tissue water) and BAI total score
